# Variation of immune cell responses in humans reveals sex-specific coordinated signaling across cell types

**DOI:** 10.1101/567784

**Authors:** Gabriela K. Fragiadakis, Zachary B. Bjornson-Hooper, Deepthi Madhireddy, Karen Sachs, Matthew H. Spitzer, Sean C. Bendall, Garry P. Nolan

## Abstract

Assessing the health and competence of the immune system is central to evaluating vaccination responses, autoimmune conditions, cancer prognosis and treatment. With an increasing number of studies examining immune dysregulation, there is a growing need for a curated reference of variation in immune parameters in healthy individuals. We used mass cytometry (CyTOF) to profile blood from 86 humans in response to 15 ex vivo immune stimuli. We present reference ranges for cell-specific immune markers and highlight differences that appear across sex and age. We identified modules of immune features that suggests there exists and underlying structure to the immune system based on signaling pathway responses across cell types. We observed increased MAPK signaling in inflammatory pathways in innate immune cells and greater overall coordination of immune cell responses in women. In contrast, men exhibited stronger STAT1 and TBK1 responses. These reference data are publicly available as a resource for immune profiling studies.

## Introduction

Immune function is critical for effective vaccine responsiveness, wound healing, and protection against infection, autoimmunity, and cancer (1). Although the ability to measure elements of the immune response has improved dramatically with advances in gene expression analysis, cytokine profiling, and flow cytometry (2-4), clinical assessment relies predominantly on the comparatively simple complete blood count as the indicator of immune health (Mayo Clinic) (5). This disparity between our true technical capabilities and the methods employed in healthcare offers the potential for dramatic improvements in assessments of immune health.

Mass cytometry has great potential as an immune monitoring tool (6-8). The ability to measure over forty proteins per single cell enables deep profiling of the immune system from a patient’s blood or tissue sample, yielding information regarding the phenotype as well as the behavior of cells, such as signaling activity (4). These and other studies have leveraged mass cytometry to assess immune differences in individuals in different clinical contexts (6, 7, 9, 10), but a reference for healthy human immune variation at steady-state is lacking.

The Cross-Species Immune Atlas presented here and in the accompanying paper by Bjornson-Hooper *et al.*† profiles blood from 86 human subjects, 88 non-human primates (rhesus macaques, cynomolgus macaques, African green monkeys), and 50 mice using mass cytometry. Each specimen was divided and exposed to 15 immune stimuli, including cytokines, growth factors and microbial products, and was then analyzed by mass cytometry using a 39-parameter immune profiling antibody panel. This project had two objectives. The first, addressed here, was to create a reference of human immune variation that can serve as a baseline in immunological studies, similar to efforts made in genomics and other fields (11), and complementary to single-cell mapping initiatives across tissues such as the Human Cell Atlas (https://www.humancellatlas.org/). The second was to determine which aspects of immune cell phenotype and signaling responses are conserved across species to guide our use of animal models in drug discovery; the accompanying paper by Bjorn-son-Hooper *et al.* describes these findings.

Great care was taken to standardize the immune stimuli and antibody panels across the five species to the largest extent possible. Technical error was minimized through the use of automation, single lots of reagents, and complete documentation practices. The data are publicly available and curated for ease of use as a reference by other researchers.

Analysis of this dataset provided a set of reference ranges for a large set of measured immune parameters and revealed coordinated sets of immune features that were grouped into modules based on correlated variation as measured across individuals. Immune modules revealed an immune system structure of coordinated signaling capacity across cell types, and enabled stratification of our donors across sex and age. We found that women had stronger inter- and intra-module correlation than men, suggesting a greater degree of immune cell signaling coordination in women, and that women demonstrated an increased signaling capacity in inflammatory signaling pathways in innate immune cells. Such differences in immune responses between sexes may inform the immune bases for observed differences in pathology such as infection susceptibility and autoimmune syndrome prevalence.

## Results

### Detailed immune profiling of healthy individuals provides a window into immune state

The development of a human immune reference required validated and well-controlled performance of assays on a large set of donors. To this end, an immune profiling panel consisting of met-al-conjugated antibodies against 23 surface antigens and 16 intracellular signaling proteins was designed, generated, and validated (Supplementary Tables 1 and 2). Intracellular signaling was investigated, as it has been informative in several immune profiling studies (7, 9, 12, 13). Analogous antibody panels were generated for human, mouse, and non-human primates to enable cross-species analyses (see Bjornson-Hooper *et al.*). Due to the potential for antibody degradation and technical variation in small-volume aliquoting, batches of antibodies were premixed and lyophilized into stable antibody cocktails for long-term storage and use (see Methods).

**Table 1:** Attributes of features contained within each module For the immune features classified into a given module, the proteins, cell types, and stimulation conditions common to the majority of features are listed. The most prevalent type of attribute is highlighted in blue.

| Module | Proteins | Cell types | Conditions |
| --- | --- | --- | --- |
| 1 | pSTAT1 | CD8 T cells, CD4 T cells, DCs, NK cells, B cells | IL-6, IFNa, IFNb, IFNg |
| 2 | pSTAT1 | Basophils, monocytes | PMA/iono, IFNa, IFNb |
| 3 | pERK1/2, pCREB, pMAPKAPK2 | CD4 T cells, CD8 T cells, B cells | PMA/iono |
| 4 | IkB | B cells, NK cells, CD4, CD8, monocytes, DCs | CD40L, TNFa, LPS, R848 |
| 5 | pP38, pERK1/2 | Monocytes, basophils | Anthrax |
| 6 | pSTAT1 | CD14 monocytes, CD16 monocytes | IFNa, IFNb, IFNg |
| 7 | pCREB, pP38, pERK1/2 | Monocytes, neutrophils, basophils | GMCSF, IL-6, R848, LPS, PMA/iono |
| 8 | pSTAT5, pSTAT6 | Monocytes, neutrophils, DCs, T cells | IL-2, GMCSF, IFNa, IFNb |
| 9 | pTBK1, pCREB, pMAPKAPK2, pP38, pERK1/2 | DCs, NK cells, monocytes | TNF |
| 10 | pSTAT4, pSTAT5, pSTAT6 | CD8, CD4, DCs, NK cells, neutrophils, monocytes | IFNa, IFNb, IL-4, IL-6 |
| 11 | pTBK1, pCREB, pMAPKAPK2, pP38, pERK1/2 | Monocytes, DCs, neutrophils | PMA/iono, LPS, R848, GMCSF |

Eighty-six healthy humans donated peripheral whole blood samples for this study (Supplementary Table 3). Samples from each volunteer were divided into 16 aliquots and were stimulated with 15 immune modulators, including cytokines, growth factors, cell type-specific agonists, and microbial antigens, or left unstimulated (Figure 1A, Supplementary Table 4). All samples from a given donor were barcoded (multiplexed), pooled, stained, and analyzed by mass cytometry simultaneously to reduce technical variability. The unstimulated condition for each donor served as an internal control to further minimize technical variability.

**Figure 1:**
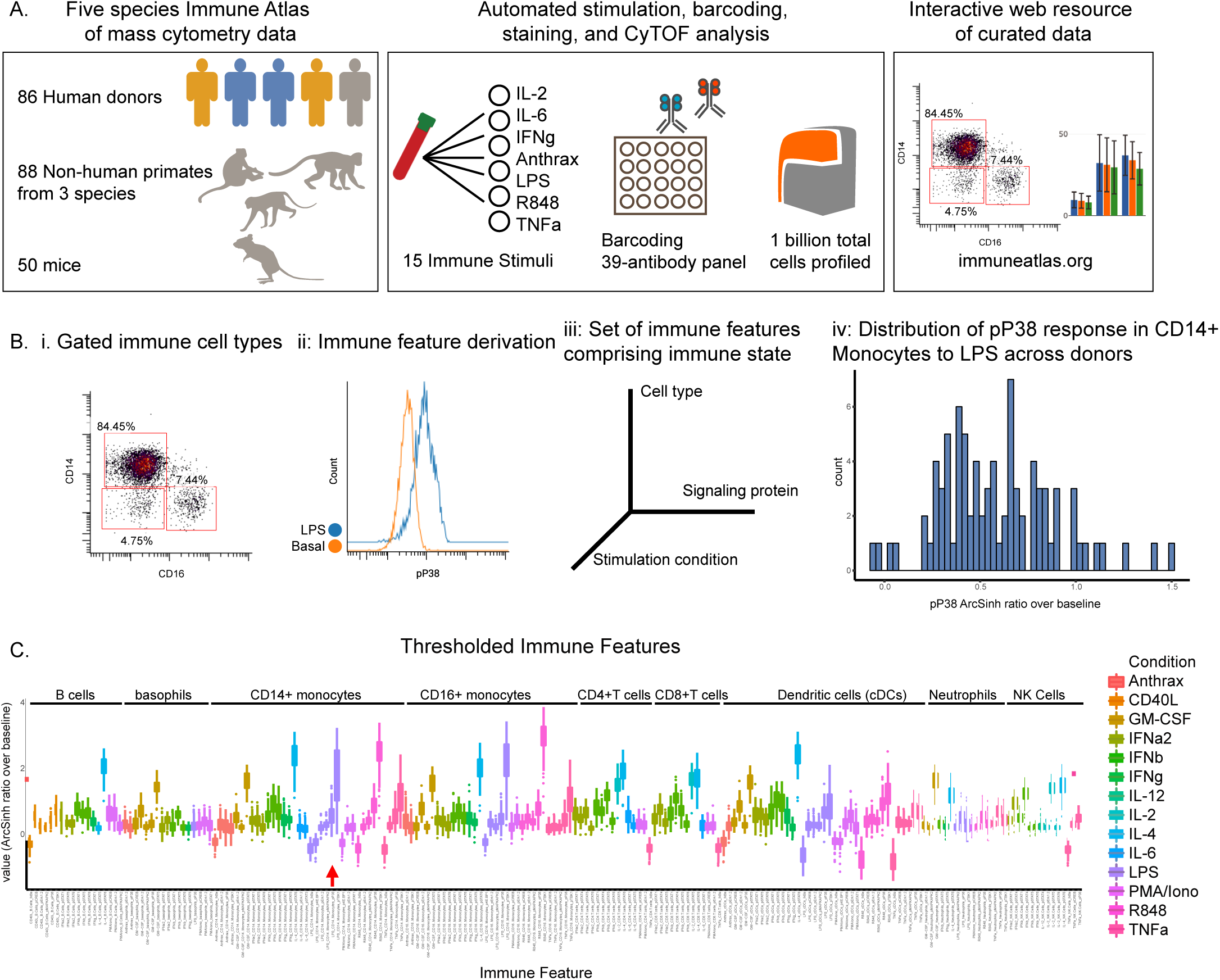
Detailed immune profiling of healthy individuals using mass cytometry (A) Generation of immune reference data set. Whole blood samples taken from 86 human donors, 88 non-human primates (rhesus macaques, cynomolgus macaques, African green monkeys), and 50 mice were divided into 16 aliquots. Aliquots were stimulated with a cytokine or microbial agent or left untreated. Samples were barcoded and stained with a 50-parameter antibody panel, and cells were analyzed by mass cytometry. The gated and curated data is available at https://flowrepository.org (accession FR-FCM-Z2ZY) and https://immuneatlas.org (B) Generation of immune features. Panel i: Cells were gated into immune cell types. Panel ii: Cell type-specific responses were calculated by subtracting the arcsinh transformed median value in the unstimulated condition from the arcsinh transformed median value in the stimulated condition (shown: p38 response to LPS in CD14+ monocytes). Panel iii: Immune state was defined as the set of immune features, and immune features were defined as the signaling response of each protein in each cell type in response to each stimulus. A total of 2,160 immune features (16 signaling proteins in 9 cell types across 15 conditions) were defined. Panel iv: The distribution of donor-specific values for the immune feature p38 response to LPS in CD14+ monocytes. (C) Box plots of reference ranges of the 199 immune features that exceeded a mean value of 0.2 arcsinh ratio. Immune features are grouped by cell type and colored by condition.

The high-dimensional nature of mass cytometry measurements combined with the large stimulation panel yielded 2,160 immune “features” from the data set. Cells were gated into nine major populations that were the focus of this investigation (Supplementary Figure 1). Note that the large number of surface markers enables other researchers to gate and analyze many additional populations than those we explore here. Immune features were defined as the level of a given signaling protein in a canonical cell type in a stimulation condition relative to the unstimulated control (Figure 1B). The set of immune features for a given donor was defined as that donor’s immune state. This analysis provided reference ranges for all 2,160 immune features based on the distribution across the eighty-six human donors. Because not every stimulus activated every signaling pathway in every cell type, we set a threshold to limit the analysis to major biological responses. This filter resulted in 199 features for further analysis (Figure 1C, Methods). This set of reference ranges of healthy immune variation is available for use in future immune profiling studies.

### Immune variation enables detection of response modules predominantly defined by signaling proteins

In order to leverage the multi-parameter profiling of the donors we used each individual in the cohort as an observed instance of a genetic or environmental perturbation of immune state. Each feature was correlated with every other feature across donors to produce a correlation map (Figures 2A, 2B) wherein hierarchical clustering placed features that were correlated near one another. To better visualize the grouping of correlated features, correlations below a stringent threshold were removed (|R| < 0.5), producing an adjacency matrix (Figure 2C). This analysis revealed highly correlated features, which were grouped into modules (within-module correlation mean R = 0.68 versus correlation between all features mean R = 0.20, Figure 2C).

**Figure 2:**
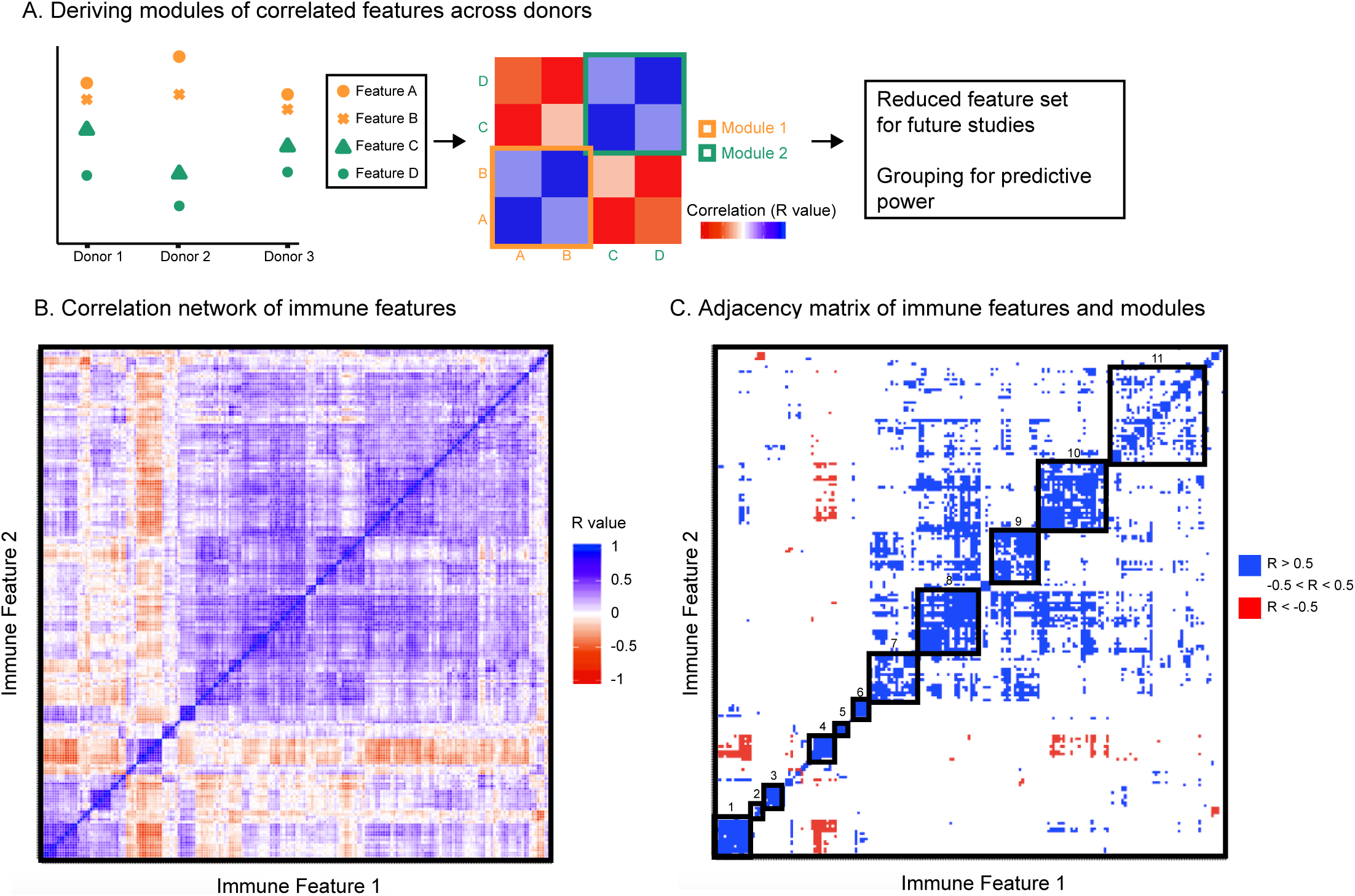
Immune variation enables detection of response modules predominantly defined by signaling proteins (A) All features were correlated with one another across donors. Highly correlated features were identified and annotated as groups of correlated features, or modules. (B) Clustered correlation heat map of the 199 features. Immune features were clustered on each axis based on similarity of correlation coefficient (R-value). Heat map is colored by R-value. (C) R-values were binned yielding an adjacency matrix. R-values from −1 to −0.5 are red, R-values from −0.5 to 0.5 are white, and R-values from 0.5 to 1 are blue. Modules were drawn based on visualized groups of highly correlated features (black boxes).

The features contained within each module were predominantly grouped by signaling protein, rather than by cell type or by stimulation condition (Table 1, Supplementary Table 5). This grouping revealed that if an individual had lower phosphorylated levels of a given signaling protein in a given cell type and condition compared to the population average, that individual was likely to also have lower levels of that phosphorylated signaling protein across other cell types and conditions. For example, if an individual had lower-than-average pSTAT1 in CD4+ T cells responding to IFN-a2, then that individual was likely to have lower levels of pSTAT1 in other cell types and conditions. This suggests a level of immune regulation and structure based on signaling pathway activity that is present across many cell types within an individual. The consistency of signaling capacity across conditions suggested that the ex vivo stimulations probed an intrinsic regulation of each signaling pathway across an individual’s cells. In contrast, modules were not organized by specific cell lineages. Whereas lymphoid and myeloid cells tended to be grouped together, several cell types shared highly correlated activity of a particular signaling pathway. Similarly, although many of the stimuli used are pleiotropic and activated several signaling pathways, modules for the most part were not organized by stimulus with the exception of Modules 3, 5, and 9. These results demonstrate that the activation of different signaling pathways elicited by a particular stimulus is not as coordinated as the activity of a given signaling pathway to different stimuli. Therefore, the immune set point in healthy individuals reveals the coordination of signaling pathway activity, which defines each person’s propensity to respond to numerous immunological stimuli.

At a practical level, these results also imply the evaluation of a smaller number of conditions may provide nearly equally meaningful information (i.e. a surrogate) with respect to general immune state. Therefore, with this dataset as a foundation, future studies using more focused diagnostic immune monitoring can be performed using a subset of proteins and conditions as a traditional flow cytometry assay (Supplementary Table 6). This is especially important in the clinical setting, where typically only flow cytometry is readily available. Surrogate markers for the majority of modules are present in this subset of proteins and conditions and their normalized values were highly correlated with derived module scores, the normalized average of the immune features within each module (Supplementary Figure 2, Methods).

### Immune structure stratifies immune responses between men and women

Having observed that measured immune responses were organized into signaling-based modules, we next sought to determine whether this organization could characterize differences in immune state between individuals. We were first interested in whether this modular structure enabled stratification by sex (Figure 3A). We performed predictive modeling, including models that allowed for input of the module assignments as a means of incorporating the higher order relationships from the data. The data were split into a training set of 50 donors and a test set of 36 donors. Four models were optimized using the training data including two regularized methods that did not use module groupings (ridge (14) and lasso (15)) and two that used the module groupings (group lasso (16) and sparse group lasso (17), see Methods). We assessed the predictive performance of each model on the test data, and found that the immune data significantly correctly classified the data by sex, the sparse group lasso model performing the best (74% correct classification on test; AUC 0.79) (Figure 3B). Notably, the model that used the internal structure of the data in addition to the immune features best defined differences in immune state between men and women.

**Figure 3:**
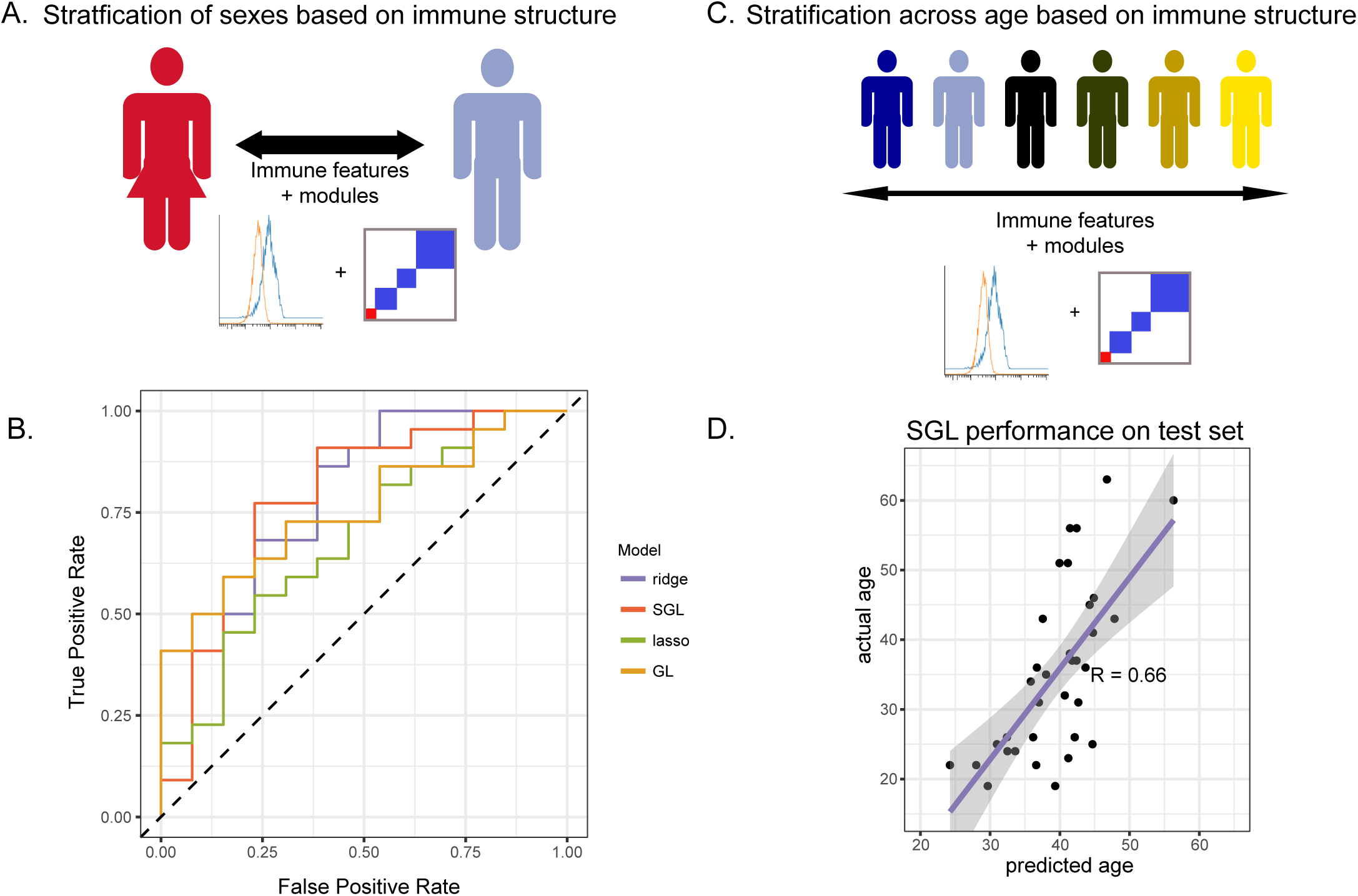
Immune modules enable improved stratification of immune responses across sex and age (A) Schematic of modeling approach for sex differences. (B) Receiver operating characteristic curve for ridge, lasso, group lasso, and sparse group lasso. Performances on test data were as follows (stated as percent correct classification, area under the curve): ridge, 68%, .78; lasso, 60%, .69; group lasso, 60%, .76; sparse group lasso, 74%, .79. (C) Schematic of modeling approach for modeling age. (D) Correlation of actual ages versus ages predicted for test set using the sparse group lasso model (R = 0.66).

The lasso model selected immune features that included signaling proteins pTBK1 and pSTAT1 (positive coefficients, higher in men) and features that included pERK1/2, pP38, and pCREB (negative coefficients, higher in women, Supplementary Table 7; see further discussion of selected features later in the manuscript). The sparse group lasso model was more inclusive and selected 35 features from modules 1, 3, 4, 5, 7, 8, 10, and 11, that included the majority of measured signaling proteins (Supplementary Table 8). To assess how much predictive information could be captured on the course-grain module level, we also trained a lasso model on personalized module scores rather than on raw immune features. The model selected module scores 1, 5, and 7, and had a classification rate of 68% on the test set (AUC 0.67). This model outperformed the lasso model and performed comparably well to our ridge model, revealing that this simplified module-based model was able to harness much of the predictive information for classification by sex (Supplementary Table 9). Taken together, these modeling results confirm that the modular structure we observed captures relevant variability in immune state, in this case revealing sex differences in immune signaling.

### Immune features and structure predict donor age

To further explore the informative value of our curated immune data, the relationship between the age of the donors and their immune responses was assessed (Figure 3C). Immune function declines with age as shown by decreased B cell and T cell function, chronic inflammation, and poorer vaccine responsiveness in elderly compared to young subjects (18-20). We examined the stratification of immune features with age by training a lasso regression model and a sparse group lasso model on the training set, using donor age as a continuous variable. The lasso regression model returned a correlation value of 0.83 on the training data and 0.64 on the test data, detecting a relationship between immune features and age (21), though the best performance was observed using the sparse group lasso which incorporates module assignments (R = 0.66, test data, Figure 3D). The lasso model selected 12 features (Supplementary Table 10); those with the largest coefficients were the pSTAT1 response in CD8 T cells to interferon, the pERK response in NK cells to IL-2, and the pMAPKAPK2 response in neutrophils to TNF stimulation. Though changes in these specific signaling pathways have not been described, these responses are consistent with previous reports of lowgrade inflammation and adaptive immuno-senescence in the elderly. Age-associated decreases in T cell function and NK cell cytokine responsiveness and increased neutrophil activity have been reported (22-24). Interestingly, though including the module assignments improved predictive performance, we could not produce a cross-validated model of age using the module scores alone, suggesting that the reductive scores do not capture sufficient predictive information in the context of age. In summary, immune features along with the relationships between features enabled the detection of both sex-associated and age-associated immune differences.

### Women have increased coordination of immune cell signaling capacity

To further explore differences in immune structure between male and female donors in our cohort, immune feature correlations were calculated separately for males and females. From the correlation networks, adjacency matrices were produced that maintained the feature order and module grouping from the full data set (Figures 4A, 4B). This analysis revealed that features from female donors were more highly correlated with one another than features were within male donors (Figure 4C, p-value = 2.2×10-16). In seven modules, within-module correlation was higher in women than in men, whereas only one module had higher correlation in men than women (Figure 4D). This suggests a higher degree of immune regulation and coordination among components of the female immune response, versus in males, where immune cell responses were relatively more independent of one another. Interestingly, when we compared the networks from younger individuals (≤ 30) to older individuals (≥ 45) there was increased correlation among immune features in older individuals (Supplementary Figure 3), perhaps reflecting the greater diversity of immune phenotypes associated with aging (21).

**Figure 4:**
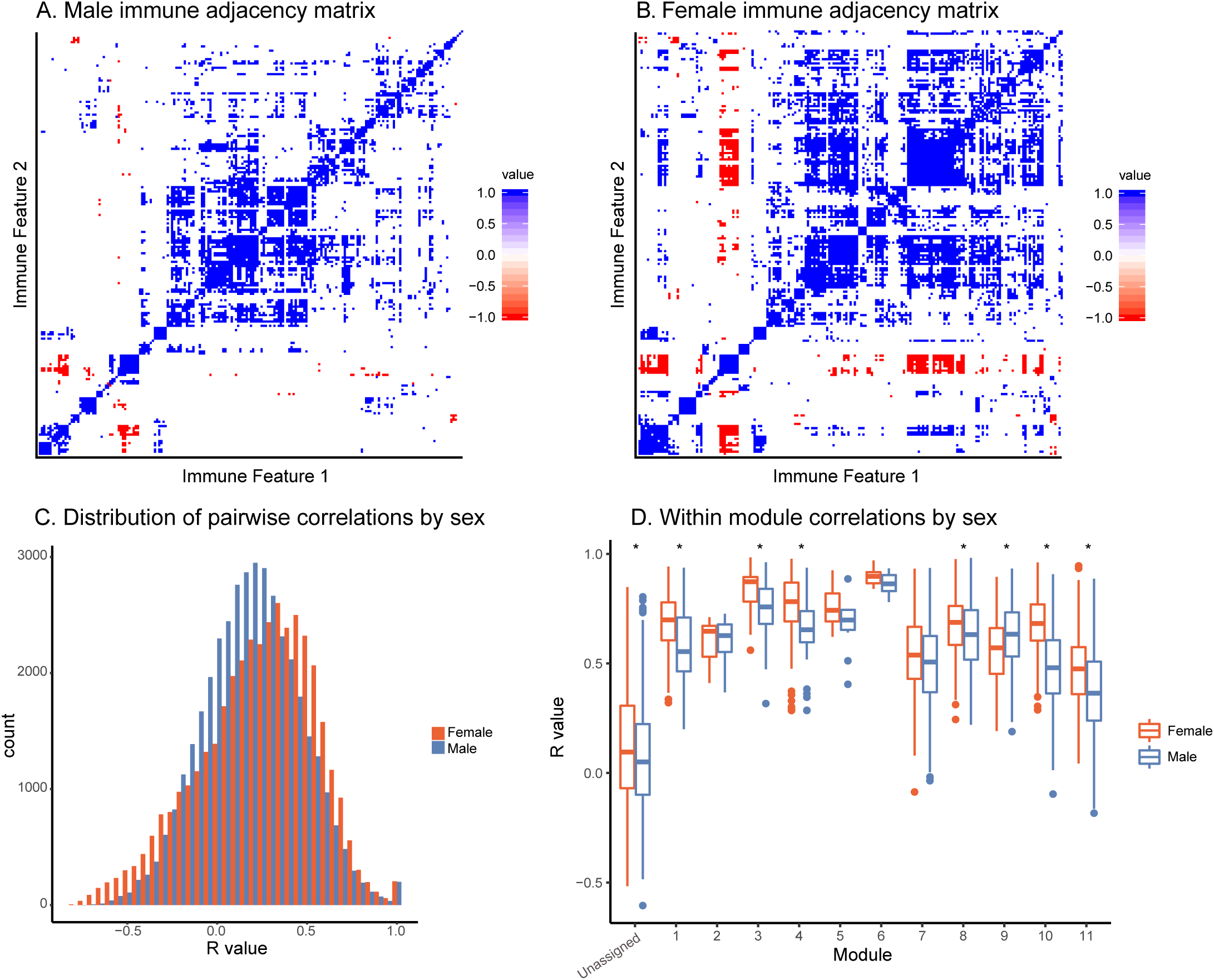
Women have increased coordination of immune cell signaling capacity (A) Adjacency matrices of 199 immune features across male donors. R-values were binned; R-values from −1 to −0.5 are red, from −0.5 to 0.5 are white, and from 0.5 to 1 are blue. Feature order was set by the clustering order on the full dataset. (B) Adjacency matrices of 199 immune features across female donors. R-values were binned; R-values from −1 to −0.5 are red, from −0.5 to 0.5 are white, and from 0.5 to 1 are blue. Feature order was set by the clustering order on the full dataset. (C) Distribution of correlation coefficients (R-values) of each pairwise feature in male donors (turquoise) compared to female donors (pink). Distributions were significantly different (p-value = 2.2×10-16, Wilcoxon sum-rank test). (D)Box-plots of correlation coefficients (R-values) within modules grouped by sex. Modules are numbered as in Figure 2C. Women (pink) had higher levels of correlation in modules 1, 3, 4, 8, 10, 11, and unassigned (Wilcoxon sum-rank test, adjusted p-value < .05). Men (turquoise) had higher levels of correlation in module 9 (Wilcoxon sum-rank test, adjusted p-value < .05).

### Men and women have distinct immune response profiles

Several studies have suggested that there are intrinsic immune differences between men and women. Women have drastically higher incidence of autoimmune disease, whereas men have poorer tolerance of infection and responses to vaccines (25, 26). These clinical differences, as well as interactions between sex-linked genes, hormones, and the immune system (27), strongly suggest that differences exist between the male and female immune systems. Module scores, averages of an individual’s normalized levels of each immune feature in a module, were significantly different between men and women in two modules (Figure 5A, Kolmogorov-Smirinov test). The module higher in men (module 1) was predominantly made up of pSTAT1 features across several lymphocyte subsets from interferon and IL-6 conditions, whereas the module higher in women (module 7) was comprised of pCREB, pP38, and pERK1/2 in innate immune cell types (Table 1). Interestingly, module 1 scores did not correlate with module 7 scores within a given sex, suggesting that on a per-individual basis having a stronger sex-based phenotype in one module does not have implications for the sex-based phenotype in the other module (Supplementary Figure 4). Other trends in module-based sex differences that did not pass our significance threshold when correcting for multiple hypothesis testing included modules 3 and 5 (higher in males) and modules 4 and 9 (higher in females) (Supplementary Figure 5).

**Figure 5:**
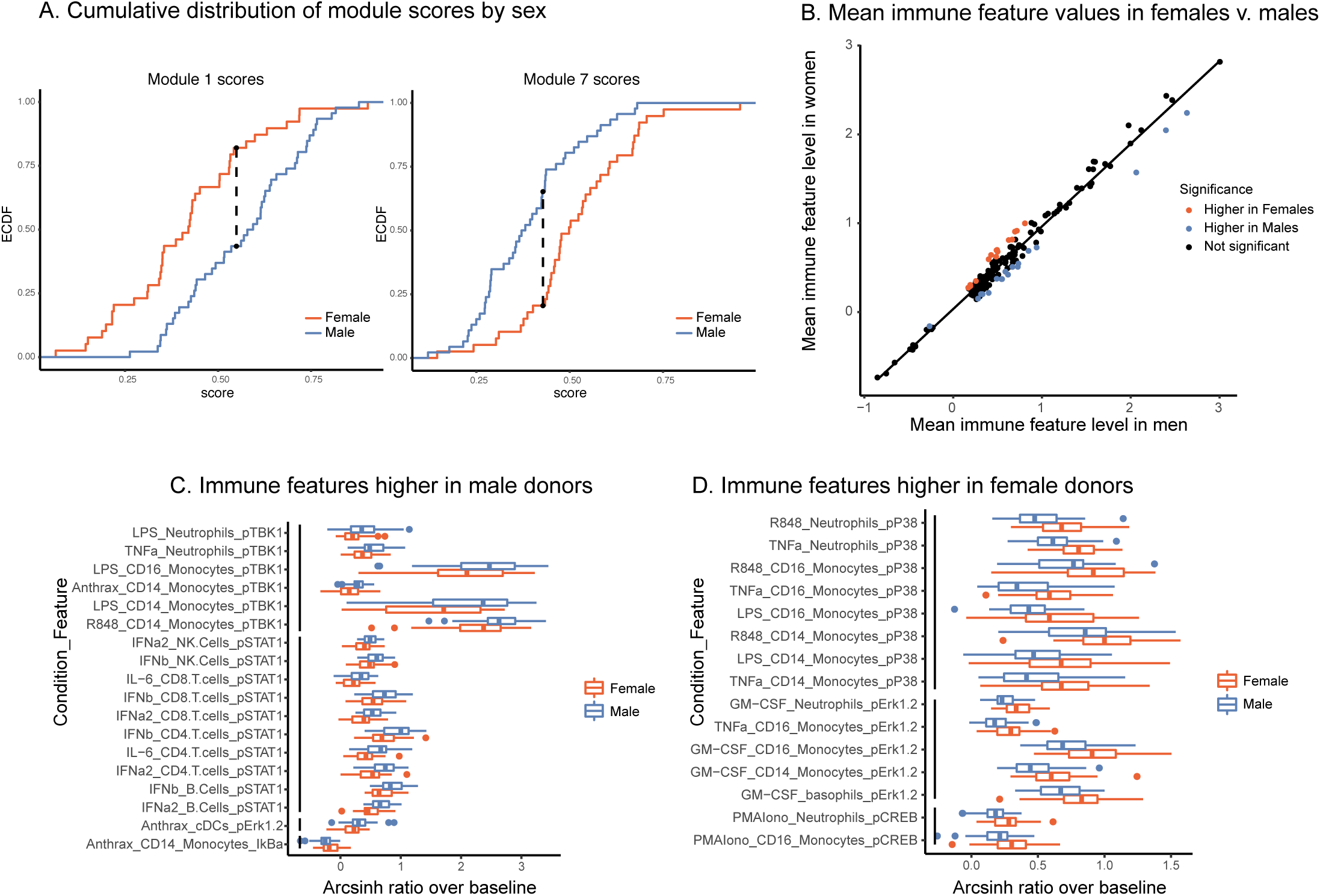
Men and women have distinct immune response profiles (A) Modules with significantly different module scores between men and women. Empirical cumulative distribution function (ECDF) shown for modules scores from module 1 (left) and module 7 (right). Dashed line shows maximum distance between distributions. Significance determined by Kolmogorov-Smirnov test, adjusted p-value < .05. (B) Mean levels of each feature in samples from female donors versus male donors. Red indicates significantly higher in females; blue, significantly higher in males; and black, not significantly different based on an FDR < 1% (SAM unpaired). (C) Boxplots of the features significantly higher in males (FDR < 1%, SAM unpaired). Features are grouped by signaling protein (black bars). Decreased IkBa plotted with male donors to reflect increased degradation of IkBa leading to higher levels of p-NFkB. (D) Boxplots of the features significantly higher in females (FDR < 1%, SAM unpaired). Features grouped by signaling protein (black bars).

To further explore this result, we performed an exhaustive assessment of all thresholded immune features in the dataset. Whereas feature medians were globally highly similar between the sexes (R = 0.985), a subset of features was significantly different between male and female donors (SAM, FDR < 0.01, Figure 5B). Differences were highly consistent by signaling protein. Of the 18 features that were significantly higher in male donors, six were pTBK1 features and 10 were pSTAT1 features across conditions and cell types (Figure 5C). Similarly, of the 15 features significantly higher in female donors, all were features involving pP38, pCREB, or pERK1/2 (Figure 5D). These were the same proteins that were selected by our lasso model of sex-differences. Interestingly, responses to TNFa, LPS, and R848 were higher in men in certain cell-type specific proteins and higher in women in others, suggesting that the response to the same stimulus can be differentially regulated across signaling pathways. In contrast, all significantly different responses to GM-CSF were exclusively higher in women than in men, whereas responses to interferons, IL-6 and Bacillus anthracis antigen were exclusively higher in men. In addition, not all pathways responsive to a given stimulus were different between sexes; for example, whereas GM-CSF stimulates both a pERK and a pSTAT5 response in monocytes, only the response to pERK was higher in women. Notably, all of the features higher in female donors were from inflammatory cell types and signaling pathways known to be involved in inflammation, consistent with higher rates of autoimmunity and enhanced responses to infection observed in women (Figure 6).

**Figure 6:**
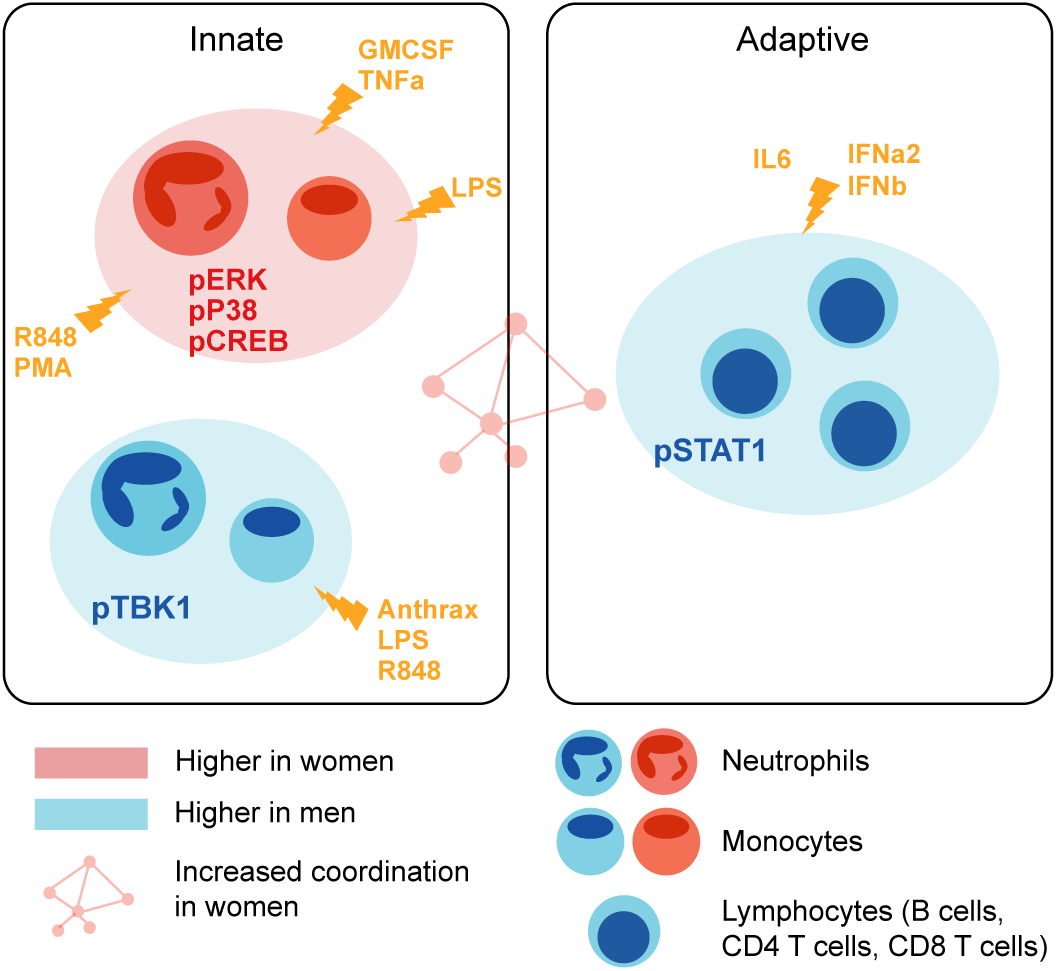
Model of immune differences between men and women Summary of immune differences detected between men and women.

## Discussion

The goals of this analysis were 1) to generate a set of mass cytometry data collected from healthy human subjects, curated with explicit immune features to be used as a reference in immune monitoring studies and 2) to leverage the multi-parameter measurements of healthy immune variation to gain insights into immune organization. This highly controlled dataset consists of immune measurements on samples from 86 individuals. Mass cytometry measurements included 16 signaling protein responses to 15 different immune modulators in nine canonical cell types. This resulted in 2,160 feature measurements for each individual, filtered to 199 features that met our threshold of responsiveness. The raw data as well as the curated and filtered immune features have been made available to the research community for use as a reference in immune monitoring studies at https://flowrepository.org (accession FR-FCM-Z2ZY) and https://immuneatlas.org.

Using each feature as a standalone measurement does not utilize the data to its full potential, as this high-dimensional dataset contains information about how the immune system is preconfigured to respond in a coordinated manner to immune stimuli. To extend the analysis beyond univariate measurements toward an understanding of the relationship between features, we leveraged the multi-parameter measurements available wherein individuals were used as instances of genetic or environmental perturbation. The merits of this type of approach have been shown with the identification of co-regulated features and module detection in gene expression data (28) and in cytometry data in specific immune modulatory contexts(12). Here we identified modules of correlated immune features across humans as a group, and men or women as sub-cohorts. The attribute common across features within a module was a shared signaling protein. This result reveals that 1) the immune system is structured in such a way that the signaling propensity of a given pathway in an individual is coordinated across cell types and across different immune perturbations and 2) that there is a degree of redundant information contained in these measurements such that future studies may perform smaller assays that can be readily performed in the clinical setting. An example of a restricted set of parameters that could be used for representative immune monitoring using traditional flow cytometry is provided in Supplementary Table 6.

In our predictive models, the elucidated immune structure informed stratification of both sex and age. Incorporating module assignments from this work, which leveraged not only the features themselves but also the relationships between immune features, enabled the detection of differences in immune responses, demonstrating that the detected internal structure captures feature relationships that vary with demographic group in humans. The modular structure improves prediction and also improves the interpretability of resulting models. This in turn suggests this framework may be useful in analyses of infection susceptibility, autoimmunity burden, and treatment response.

Analysis of this dataset allowed for a detailed characterization of differences between the male and female immune systems. Significantly higher responses in male donors than in females were predominantly detected in pSTAT1 in lymphocytes and pTBK1 in myeloid subsets, whereas female donors had higher levels of pERK1/2, pCREB, and pP38 features in monocytes and neutrophils (Figure 6). The female immune system has been shown to have higher inflammatory phenotypes than the male immune system both clinically and at the cellular level (29), which is consistent with our findings of increased MAP-kinase signaling in innate immune inflammatory cells in women compared to men. In contrast, neutrophils and monocytes were more abundant in males than in females (Supplementary Figure 6). Perhaps this disparity is the reason that inflammatory pathways within monocytes and neutrophils have a greater response capacity in women. Notably, most immune profiling studies using flow cytometry examine only cell frequencies rather than cell-specific signaling pathways responses, and therefore do not capture this larger set of differences. Interestingly, males have worse outcomes in a multitude of microbial infections (26). It was therefore surprising that pSTAT1 and pTBK1 capacity was higher in men than women; this could be a compensatory mechanism to counter the fact that males have lower interferon alpha production in response to inflammatory stimuli and therefore require more sensitive signaling responses (30).

Our analysis also revealed more nuanced aspects of sex-specific immune regulation. Women had a greater degree of correlation of responses across immune features than men, which may point to greater coordination of immune responses and structure. It is unclear why immune cell signaling responses in males are more independent of one another, although it may reflect a more defined immune set point in women. Future work will be required to examine the nature of this sex-specific coordination.

Although it is clear that immune structure can reveal differences in demographic parameters, here sex and age, its utility has yet to be shown in instances of immune pathology. Because the effects of pathology may exceed those of either sex or age on immune health, it is likely that exploring higher-order interactions between immune features will be informative in clinical studies. It will be interesting to see whether these relationships tend to hold or break down in those contexts. The dataset reported here includes only data collected from a single blood draw from each donor, as opposed to longitudinal draws from the same donor. Longitudinal studies are needed to determine how stable the immune state of a healthy donor is and how consistent it is in its response to an immune modulator. Such longitudinal studies will indicate, for example, whether there is an optimal time for introduction of a perturbation such as vaccination, or a time when an individual is more or less susceptible to infection. Future work should be aimed at integrating different complementary sources of immune information, such as gene expression and cytokine analysis, to provide a richer picture of immune state. This work and the accompanying manuscript by Bjornson-Hooper *et al.* provide the first analyses of the Cross-Species Immune Atlas dataset. This curated resource as well as these initial analyses will enable future human immune monitoring studies as well as more rational pre-clinical studies.

## Methods

The methods for reagent development and sample processing are those used for the Cross-Species Immune Atlas project and are redundantly stated here and in Bjornson-Hooper *et al.*

### Data generation

#### Stimuli

Reagents used for stimulations are listed in Supplementary Table 4. Stimuli were tested in whole blood over a range of concentrations to select the working concentration. All reagents were diluted such that the same volume of each achieved the desired level of stimulation. Stimulation reagents were then aliquoted into single-use plates and stored at −80 °C with the exception of those indicated in Supplementary Table 4, which were dispensed at time of use due to storage requirements. All cytokines were tested for endotoxin by the LAL method and verified to contain an amount less than that detectable by our phospho-flow assays (approximately 10 pg/ ml). The LPS used was prepared by phenol-water extraction and contained small amounts of other bacterial components that activate TLR2. Gamma-inactivated vegetative Bacillus anthracis Ames (ANG-BACI008-VE) was obtained from the Department of Defense Critical Reagents Program through the NIH Biodefense and Emerging Infections Research Resources Repository, NIAID, NIH.

#### Blood

Venous human blood was obtained from the Stanford Blood Center and from AllCells (exempt, non-human subjects research) and from volunteers from the Stanford community under an IRB-approved protocol.

#### Antibodies

Purified antibodies were purchased and conjugated in-house using DVS/Fluidigim MaxPar X8 metal conjugation kits. All antibodies were titrated for optimal signal-to-noise ratio, which was confirmed in at least two different individuals per species (three humans, two cynomolgus macaques, two rhesus macaques, three mice). All conjugations and titrations were well-documented, and records are available in the experiment data repository. Antibodies were lyophilized into LyoSpheres by BioLyph with excipient B144 as 4x cocktails. CyTOF antibody LyoSpheres were stress-tested for over one year and found to have no significant change in staining (not shown).

#### Stimulation and staining

Stimulations and staining were carried out on an automation platform consisting of an Agilent Bravo pipetting robot, Agilent BenchBot robotic arm, Peak KiNeDx robotic arm, Thermo Cytomat C2 incubator, BioTek ELx405-UVSD aspirator/dispenser, BioTek MultiFlo FX four-reagent dispenser, Q.Instruments microplate shakers, Velocity11 VSpin centrifuges, and a custom chilling system contained in a negative-pressure biosafety enclosure. The VWorks robotic programs and log files are available upon request.

Whole blood was stimulated by mixing with stimulus reagent and incubating in a humidified 37 °C, 5% CO2 incubator for 15 minutes. Blood was fixed for 10 minutes at room temperature with 1.6% paraformaldehyde (PFA, Electron Microscopy Sciences) and lysed with 0.1% Triton-X100 in PBS for 30 minutes at room temperature (31). Cells were washed twice with PBS, then each of the 16 conditions for each donor was barcoded as previously described (32). Briefly, cells were slightly permeabilized with 0.02% saponin, then stained with unique combinations of functionalized, stable palladium isotopes. Each stimulation plate contained samples from 6 donors stimulated under 16 conditions. After stimulation, each plate was reduced to 6 wells, each containing the 16 conditions for one donor. Cells were washed once with staining media (CSM: 0.2% BSA in PBS with 0.02% sodium azide), blocked with human TruStain FcX block (Biolegend) for 10 minutes at room temperature with shaking, then stained with rehydrated extracellular LyoSpheres for 30 minutes at room temperature with shaking in a final volume of 240 µl. Cells were washed once with CSM, then permeabilized in >90% methanol at 4 °C for 20 minutes. Cells were washed four times, then stained with intracellular lyospheres for 60 minutes at room temperature with shaking. Cells were washed once, then placed into 1.6% PFA and 0.1 µM natural iridium intercalator (Fluidigm) in PBS at 4 °C until acquisition on a CyTOF. With few exceptions, cells were acquired within seven days of staining. From prior validation experiments, this amount of time imparted no significant effect on staining.

#### Acquisition

Prior to analysis on the CyTOF, cells were washed twice with water. Samples were acquired on a single DVS/Fluidigm CyTOF 2 fitted with a Super Sampler sample introduction system (Victorian Airship & Scientific Apparatus). Machine QC reports were run on the CyTOF between every barcoded set. Prior to data acquisition, the instrument was demonstrated to have Tb159 dual counts greater than 1,000,000 and oxidation less than 3%; if the instrument failed those criteria, it was cleaned, tuned or repaired as necessary. Approximately 4,800,000 events were acquired per sample.

### Data Processing

#### Data normalization, debarcoding, and manual gating

Data were normalized and debarcoded using the data normalization software (33) and the single cell debarcoder tool described in(34) described previously. Data were analyzed on https://immuneatlas.org.

#### Derivation of immune features

Immune features were defined as follows. A standard data transformation was applied (transformed value = arcsinh(value/5)), and then the median values of each signaling protein (n = 15) were extracted per gated cell type (n = 9) for each condition. The medians from the unstimulated condition were subtracted from the medians from each stimulation condition (n = 16). The resulting value was defined as the immune feature measurement. This derivation resulted in 2,160 immune features (16*9*15).

In the interest of removing features that were not a result of a response to the stimulus, a threshold was set empirically at a mean immune feature value of 0.2. This threshold was chosen by looking at T cell responses versus monocyte responses to LPS (T cells do not have the LPS receptor, whereas monocytes do).

### Statistical modeling

#### Sex differences in immune features

Significance of immune features between sexes was assessed using Significance Analysis of Microarrays (SAM)(35) using a false discovery rate of less than 1%. Analysis was performed in R using the samr library. Sex differences in module scores were assessed using the Kolmogorov-Smirinov test, with a Benjamini-Hochberg correction for the twelve tests performed (11 modules + unassigned).

#### Regression models

Models predicting age and gender based on immune features were optimized and evaluated using the R libraries glmnet for lasso, ridge, and elastic net; SGL for sparse group lasso; and grplasso for group lasso. The training set was initially sampled at random and kept consistent throughout for each outcome variable. Model performance was assessed on the samples not selected for the training set. For elastic net, the alpha parameter was tuned using k-fold cross-validation on three separate iterations of assigning k. This revealed that ridge (alpha = 0) had the lowest cross-validation error in each iteration (Supplementary Figure 7).

#### Module derivation, visualizations, and scores

Modules were drawn and visualized, with the ability to extract feature groupings, using an R interface (in Shiny) written for this project. Remaining visualizations shown were produced in R using the ggplot2 library. Module scores were calculated based on the average of the normalized values of the features in each module.

## Disclaimer and Acknowledgements

The research discussed in this article was supported in part by the U.S. Food and Drug Administration (Contract No. HHS-F223201210194C). This article reflects the views of the authors and should not be construed to represent the U.S. Food and Drug Administration’s views or policies. G.K.F. was supported by the Stanford Bio-X graduate research fellowship and NIH grant T32GM007276. Z.B.B.H. was supported by NIH grant T32GM007276. M.H.S. was supported by NIH grant DP5OD023056. Additional support was provided by NIH awards 5R01CA18496804, 5R25CA18099304, 1R01GM10983604, 5UH2AR06767603, 1R01NS08953304 and R01HL120724.

## Author contributions

G.K.F. wrote the manuscript, contributed to data generation, conceived of and performed the data analysis. Z.B.B.H. generated the data and edited the manuscript. D.M. contributed to data generation and reagent optimization. K.S. advised data analysis and edited the manuscript. M.H.S. contributed to data generation, advised data analysis, and edited the manuscript. S.C.B. provided advising and edited the manuscript. G.P.N. advised the study and edited the manuscript.

## References

† Zachary B Bjornson-Hooper, Gabriela K Fragiadakis, Matthew H Spitzer, Deepthi Madhireddy, Dave McIlwain, Garry P Nolan. A comprehensive atlas of immunological differences between humans, mice and non-human primates. Submitted to Biorxiv (DOI pending), 2019.

2. L. Zitvogel, A. Tesniere, G. Kroemer, Cancer despite immunosurveillance: immunoselection and immunosubversion. Nature reviews. Immunology 6, 715–727 (2006).

3. Z. Wang, M. Gerstein, M. Snyder, RNA-Seq: a revolutionary tool for transcriptomics. Nature reviews. Genetics 10, 57–63 (2009).

4. H. E. Lynch et al., Development and implementation of a proficiency testing program for Luminex bead-based cytokine assays. Journal of immunological methods 409, 62–71 (2014).

5. S. C. Bendall et al., Single-cell mass cytometry of differential immune and drug responses across a human hematopoietic continuum. Science 332, 687–696 (2011). (Mayo Clinic).

6. B. Gaudilliere et al., Clinical recovery from surgery correlates with single-cell immune signatures. Science translational medicine 6, 255ra131 (2014).

7. G. K. Fragiadakis et al., Patient-specific Immune States before Surgery Are Strong Correlates of Surgical Recovery. Anesthesiology 123, 1241– 1255 (2015).

8. M. H. Spitzer et al., IMMUNOLOGY. An interactive reference framework for modeling a dynamic immune system. Science 349, 1259425 (2015).

9. W. E. O’Gorman et al., Single-cell systems-level analysis of human Toll-like receptor activation defines a chemokine signature in patients with systemic lupus erythematosus. The Journal of allergy and clinical immunology 136, 1326–1336 (2015).

10. P. Brodin et al., Variation in the human immune system is largely driven by non-heritable influences. Cell 160, 37–47 (2015).

11. C. Genomes Project et al., A map of human genome variation from population-scale sequencing. Nature 467, 1061–1073 (2010).

12. A. N. Hotson et al., Coordinate actions of innate immune responses oppose those of the adaptive immune system during Salmonella infection of mice. Science signaling 9, ra4 (2016).

13. J. M. Irish et al., Single cell profiling of potentiated phospho-protein networks in cancer cells. Cell 118, 217–228 (2004).

14. A. E. K. Hoerl, Robert W., Ridge regression: Biased estimation for nonorthogonal problems. Technometrics 12.1, 55–67 (1970).

15. R. Tibshirani, Regression Shrinkage and Selection via the lasso. Journal of the Royal Statistical Society. Series B (methodological) 58, 267–288 (1996).

16. M. L. Yuan, Yi Model Selection and Estimation in Regression with Grouped Variables. Journal of the Royal Statistical Society. Series B (statistical Methodology) 68, 49–67 (2006).

17. N. F. Simon, J; Hastie T; Tibshirani, R A Sparse-Group Lasso. Journal of Computational and Graphical Statistics 22, 231–245 (2013).

18. L. Haynes, A. C. Maue, Effects of aging on T cell function. Current opinion in immunology 21, 414–417 (2009).

19. A. C. Shaw, S. Joshi, H. Greenwood, A. Panda, J. M. Lord, Aging of the innate immune system. Current opinion in immunology 22, 507–513 (2010).

20. P. V. Targonski, R. M. Jacobson, G. A. Poland, Immunosenescence: role and measurement in influenza vaccine response among the elderly. Vaccine 25, 3066–3069 (2007).

21. S. S. Shen-Orr et al., Defective Signaling in the JAK-STAT Pathway Tracks with Chronic Inflammation and Cardiovascular Risk in Aging Humans. Cell Syst 3, 374–384 e374 (2016).

22. D. Zhang et al., Neutrophil ageing is regulated by the microbiome. Nature 525, 528–532 (2015).

23. M. Czesnikiewicz-Guzik et al., T cell subset-specific susceptibility to aging. Clinical immunology 127, 107–118 (2008).

24. E. Mariani et al., Different IL-8 production by T and NK lymphocytes in elderly subjects. Mechanisms of ageing and development 122, 1383–1395 (2001).

25. P. Invernizzi, S. Pasini, C. Selmi, M. E. Gershwin, M. Podda, Female predominance and X chromosome defects in autoimmune diseases. Journal of autoimmunity 33, 12–16 (2009).

26. S. L. Klein, The effects of hormones on sex differences in infection: from genes to behavior. Neuroscience and biobehavioral reviews 24, 627–638 (2000).

27. E. N. Fish, The X-files in immunity: sex-based differences predispose immune responses. Nature reviews. Immunology 8, 737–744 (2008).

28. E. Segal et al., Module networks: identifying regulatory modules and their condition-specific regulators from gene expression data. Nature genetics 34, 166–176 (2003).

29. D. Furman et al., Systems analysis of sex differences reveals an immunosuppressive role for testosterone in the response to influenza vaccination. Proceedings of the National Academy of Sciences of the United States of America 111, 869–874 (2014).

30. A. Meier et al., Sex differences in the Toll-like receptor-mediated response of plasmacytoid dendritic cells to HIV-1. Nature medicine 15, 955–959 (2009).

31. S. Chow et al., Whole blood fixation and permeabilization protocol with red blood cell lysis for flow cytometry of intracellular phosphorylated epitopes in leukocyte subpopulations. Cytometry. Part A: the journal of the International Society for Analytical Cytology 67, 4–17 (2005).

32. G. K. Behbehani et al., Transient partial permeabilization with saponin enables cellular barcoding prior to surface marker staining. Cytometry. Part A: the journal of the International Society for Analytical Cytology 85, 1011–1019 (2014).

33. R. Finck et al., Normalization of mass cytometry data with bead standards. Cytometry. Part A: the journal of the International Society for Analytical Cytology 83, 483–494 (2013).

34. E. R. Zunder et al., Palladium-based mass tag cell barcoding with a doublet-filtering scheme and single-cell deconvolution algorithm. Nature protocols 10, 316–333 (2015).

35. V. G. Tusher, R. Tibshirani, G. Chu, Significance analysis of microarrays applied to the ionizing radiation response. Proceedings of the National Academy of Sciences of the United States of America 98, 5116–5121 (2001).

