## Supplementary_figures_and_tables for "Variation of immune cell responses in humans reveals sex-specific coordinated signaling across cell types"

Supplementary Data

Supplementary Table 1: Extracellular antibodies

| Antigen | Clone | Label | Vendor and Catalog | Staining Concentration (µg/ml) |
|---|---|---|---|---|
| CD45 | HI30 | 115 | Biolegend 304002 | 3.25 |
| CD235a | HIR2 | 113 | Biolegend 306602 | 0.67 |
| CD61 | VI-PL2 | 140 | Biolegend 336402 | 5.37 |
| CD66 | YTH71.3 | 158 | Pierce MA1-36189 | 3.41 |
| CD19 | J3-119 | 173 | Beckman Coulter IM1313 | 1.79 |
| IgM | G20-127 | 174 | BD 555780 | 2.68 |
| HLA-DR | Immu357 | 176 | Beckman Coulter Immu357 | 0.34 |
| CD1c | AD5-8E7 | 162 | Miltenyi (Custom) | 1.34 |
| BDCA3 | 1A4 | 164 | BD 559780 | 4.31 |
| CD123 | 7G3 | 148 | BD 554527 | 1.34 |
| CCR7 | 150503 | 171 | R&D MAB197-100 | 7.01 |
| CD3 | SP34.2 | 157 | BD 551916 | 2.68 |
| CD4* | OKT4 | 156 | Biolegend 317402 | 1.25 |
| CD8 | RPA-T8 | 155 | Biolegend 301002 | 1.41 |
| CD45RA | HI100 | 166 | Biolegend 304102 | 2.68 |
| CD11b | ICRF44 | 153 | Biolegend 301302 | 7.16 |
| CD11c | Bu15 | 143 | Biolegend 337202 | 1.34 |
| CD14 | M5E2 | 151 | Biolegend 301802 | 10.73 |
| CD16 | 3G8 | 159 | Biolegend 302033 | 2.68 |
| CD33 | AC104.3E3 | 142 | Miltenyi (Custom) | 3.58 |
| CD56 | NCAM16.2 | 175 | BD 559043 | 1.79 |
| CD7 | M-T701 | 141 | BD 555359 | 5.37 |
| CD161 | HP-G310 | 168 | Biolegend 339902 | 5.37 |

*At time of rehydration, surface cocktail was supplemented with 8.64 µl of CD4 #201 (OKT4) per 12 LyoSpheres to increase signal.

Supplementary Table 2: Intracellular antibodies

| Antigen | Label | Clone | Straining Concentration (μg/ml) |
|---|---|---|---|
| STAT1 pY701 | 147 | 4a | 5.82 |
| STAT3 pY705 | 139 | 4 | 5.82 |
| STAT4 pY693 | 170 | 38 | 5.66 |
| STAT5 pY694 | 149 | 46 | 5.37 |
| STAT6 pY691 | 165 | 18 | 5.38 |
| Ki67 | 169 | SolA15 | 3.36 |
| Erk1/2 pT202/Y204 | 152 | D13.14.4E | 16.44 |
| MAPKAPK2 pT334 | 144 | 27B7 | 2.29 |
| CREB pS133 | 145 | 87G3 | 7.24 |
| IkBa amino-terminal | 163 | L35A5 | 3.79 |
| TBK1/NAK pS172 | 161 | D52C2 | 10.78 |
| S6 pS235/236 | 150 | 2F9 | 13.15 |
| Zap70/Syk pY319/Y352 | 160 | 17a | 2.14 |
| 4E-BP1 pT37/46 | 172 | 236B4 | 3.13 |
| PLCγ2 pY759 | 146 | K86-689.37 | 2.18 |
| P38 pT180/Y182 | 154 | 36/p38 | 3.56 |
| FoxP3 | 167 | PCH101 | 10.55 |

Supplementary Table 3: Demographics

| | |
|---|---|
| **Total donors** | N = 86 |
| **Sex** | Women: N = 86<br>Men: N = 39<br>Undisclosed: N = 1 |
| **Weight** | Mean = 77.0 kg<br>SD = 18.0 kg |
| **Height** | Mean = 172.0 in<br>SD = 9.73 in |
| **Ethnicity** | Caucasian: N = 51<br>Asian: N = 10<br>African American: N = 8<br>Hispanic: N = 15<br>Native American: N = 1<br>Undisclosed: N = 1 |
| **Age** | Mean = 36.2 yrs<br>SD = 11.8 yrs |

Supplementary Table 4

Table of stimuli

| **Stimulus** | **Used In** | **Produced In** | **Lot** | **Working Concentration** |
|---|---|---|---|---|
| GM-CSF | Human | E. coli | 3112711 | 100 ng/ml |
| IFNα2 | Human | E. coli | 5962 | 150 ng/ml |
| LPS | Human | E. coli O111:B4 | LEB-36-01 | 1 μg/ml |
| IL-6 | Human | E. coli | 3103810 | 500 ng/ml |
| Resiquimod (R848) | Human | N/A | 848-35-14 | 10 μg/ml |
| IFNγ | Human | E. coli | RAX1814011 | 330 ng/ml |
| TNFα | Human | E. coli | DDHB0113062 | 100 ng/ml |
| IFNβ* | Human | CHO | 5886 | 5 ng/ml |
| CD40L soluble dimer ("MegaCD40L") | Human | CHO | 05281412, 03041401 pooled | 125 ng/ml |
| PMA and ionomycin* | Human | N/A | E13495-116 | 0.081 μM PMA and 1.34 μM iono |
| IL-12 | Human | CHO | 0210596, 0707596-2 pooled | 400 ng/ml |
| IL-4 | Human | E. coli | AG1314021 | 125 ng/ml |
| IL-2 | Human | E. coli | 101312, 041412 pooled | 2 μg/ml |
| *Gamma-inactivated vegetative *Bacillus anthracis* Ames | Human | | AGD0001331 | 400,000 CFU/ml |
| *Zaïre Ebolavirus-like particles | Human | 293T cells | | |

* Stimulation reagents were aliquoted into single-use stimuli plates and stored at -80 °C with the exception of those marked, which were dispensed at time of use.

Supplementary Table 5: Module assignments

| | Condition_Feature | Module |
|---|---|---|
| 1 | IFNa2_B.Cells_pSTAT1 | 1 |
| 2 | IFNa2_CD4.T.cells_pSTAT1 | 1 |
| 3 | IFNa2_CD8.T.cells_pSTAT1 | 1 |
| 4 | IFNa2_cDCs_pSTAT1 | 1 |
| 5 | IFNa2_NK.Cells_pSTAT1 | 1 |
| 6 | IFNb_B.Cells_pSTAT1 | 1 |
| 7 | IFNb_CD4.T.cells_pSTAT1 | 1 |
| 8 | IFNb_CD8.T.cells_pSTAT1 | 1 |
| 9 | IFNb_cDCs_pSTAT1 | 1 |
| 10 | IFNb_NK.Cells_pSTAT1 | 1 |
| 11 | IFNg_B.Cells_pSTAT1 | 1 |
| 12 | IFNg_cDCs_pSTAT1 | 1 |
| 13 | IL.6_CD4.T.cells_pSTAT1 | 1 |
| 14 | IL.6_CD8.T.cells_pSTAT1 | 1 |
| 15 | IFNa2_basophils_pSTAT1 | 2 |
| 16 | IFNb_basophils_pSTAT1 | 2 |
| 17 | PMAIono_basophils_pSTAT1 | 2 |
| 18 | PMAIono_CD14..Monocytes_pSTAT1 | 2 |
| 19 | PMAIono_B.Cells_pCREB | 3 |
| 20 | PMAIono_B.Cells_pErk1.2 | 3 |
| 21 | PMAIono_B.Cells_pMAPKAPK2 | 3 |
| 22 | PMAIono_basophils_pErk1.2 | 3 |
| 23 | PMAIono_CD4.T.cells_pCREB | 3 |
| 24 | PMAIono_CD4.T.cells_pErk1.2 | 3 |
| 25 | PMAIono_CD8.T.cells_pCREB | 3 |
| 26 | PMAIono_CD8.T.cells_pErk1.2 | 3 |
| 27 | CD40L_B.Cells_IkBa | 4 |
| 28 | LPS_CD14..Monocytes_IkBa | 4 |
| 29 | LPS_cDCs_IkBa | 4 |
| 30 | R848_CD14..Monocytes_IkBa | 4 |
| 31 | R848_cDCs_IkBa | 4 |
| 32 | TNFa_CD14..Monocytes_IkBa | 4 |
| 33 | TNFa_CD4.T.cells_IkBa | 4 |
| 34 | TNFa_CD8.T.cells_IkBa | 4 |
| 35 | TNFa_cDCs_IkBa | 4 |
| 36 | TNFa_NK.Cells_IkBa | 4 |

| | | |
|---|---|---|
| 37 | Anthrax_basophils_pErk1.2 | 5 |
| 38 | Anthrax_basophils_pP38 | 5 |
| 39 | Anthrax_CD14..Monocytes_pErk1.2 | 5 |
| 40 | Anthrax_CD14..Monocytes_pP38 | 5 |
| 41 | Anthrax_CD16..Monocytes_pErk1.2 | 5 |
| 42 | IFNa2_CD14..Monocytes_pSTAT1 | 6 |
| 43 | IFNa2_CD16..Monocytes_pSTAT1 | 6 |
| 44 | IFNb_CD14..Monocytes_pSTAT1 | 6 |
| 45 | IFNb_CD16..Monocytes_pSTAT1 | 6 |
| 46 | IFNg_CD14..Monocytes_pSTAT1 | 6 |
| 47 | IFNg_CD16..Monocytes_pSTAT1 | 6 |
| 48 | GM.CSF_basophils_pCREB | 7 |
| 49 | GM.CSF_basophils_pErk1.2 | 7 |
| 50 | GM.CSF_basophils_pP38 | 7 |
| 51 | GM.CSF_CD14..Monocytes_pErk1.2 | 7 |
| 52 | GM.CSF_CD16..Monocytes_pCREB | 7 |
| 53 | GM.CSF_CD16..Monocytes_pErk1.2 | 7 |
| 54 | GM.CSF_Neutrophils_pErk1.2 | 7 |
| 55 | IL.6_CD14..Monocytes_pErk1.2 | 7 |
| 56 | LPS_CD14..Monocytes_pErk1.2 | 7 |
| 57 | LPS_CD14..Monocytes_pP38 | 7 |
| 58 | LPS_CD16..Monocytes_pP38 | 7 |
| 59 | PMAIono_CD14..Monocytes_pErk1.2 | 7 |
| 60 | R848_CD14..Monocytes_pErk1.2 | 7 |
| 61 | R848_CD14..Monocytes_pP38 | 7 |
| 62 | R848_CD16..Monocytes_pCREB | 7 |
| 63 | R848_CD16..Monocytes_pP38 | 7 |
| 64 | R848_Neutrophils_pP38 | 7 |
| 65 | TNFa_Neutrophils_pP38 | 7 |
| 66 | GM.CSF_CD14..Monocytes_pSTAT5 | 8 |
| 67 | GM.CSF_CD16..Monocytes_pSTAT5 | 8 |
| 68 | GM.CSF_cDCs_pSTAT5 | 8 |
| 69 | GM.CSF_Neutrophils_pSTAT5 | 8 |
| 70 | IFNa2_B.Cells_pSTAT5 | 8 |
| 71 | IFNa2_B.Cells_pSTAT6 | 8 |
| 72 | IFNa2_CD14..Monocytes_pSTAT5 | 8 |
| 73 | IFNa2_CD14..Monocytes_pSTAT6 | 8 |
| 74 | IFNa2_CD16..Monocytes_pSTAT5 | 8 |
| 75 | IFNa2_CD16..Monocytes_pSTAT6 | 8 |

| | | |
|---|---|---|
| 76 | IFNb_B.Cells_pSTAT5 | 8 |
| 77 | IFNb_B.Cells_pSTAT6 | 8 |
| 78 | IFNb_CD14..Monocytes_pSTAT5 | 8 |
| 79 | IFNb_CD14..Monocytes_pSTAT6 | 8 |
| 80 | IFNb_CD16..Monocytes_pSTAT5 | 8 |
| 81 | IFNb_CD16..Monocytes_pSTAT6 | 8 |
| 82 | IFNb_cDCs_pSTAT5 | 8 |
| 83 | IFNb_cDCs_pSTAT6 | 8 |
| 84 | IFNb_Neutrophils_pSTAT5 | 8 |
| 85 | IFNg_CD14..Monocytes_pSTAT5 | 8 |
| 86 | IFNg_CD16..Monocytes_pSTAT5 | 8 |
| 87 | IFNg_Neutrophils_pSTAT5 | 8 |
| 88 | IL.2_CD4.T.cells_pSTAT5 | 8 |
| 89 | IL.2_CD8.T.cells_pSTAT5 | 8 |
| 90 | IL.2_NK.Cells_pSTAT5 | 8 |
| 91 | GM.CSF_cDCs_pCREB | 9 |
| 92 | GM.CSF_cDCs_pErk1.2 | 9 |
| 93 | LPS_cDCs_pCREB | 9 |
| 94 | R848_cDCs_pCREB | 9 |
| 95 | TNFa_CD14..Monocytes_pMAPKAPK2 | 9 |
| 96 | TNFa_CD14..Monocytes_pP38 | 9 |
| 97 | TNFa_CD14..Monocytes_pTBK1 | 9 |
| 98 | TNFa_CD16..Monocytes_pCREB | 9 |
| 99 | TNFa_CD16..Monocytes_pErk1.2 | 9 |
| 100 | TNFa_CD16..Monocytes_pMAPKAPK2 | 9 |
| 101 | TNFa_CD16..Monocytes_pP38 | 9 |
| 102 | TNFa_CD16..Monocytes_pTBK1 | 9 |
| 103 | TNFa_cDCs_pCREB | 9 |
| 104 | TNFa_cDCs_pErk1.2 | 9 |
| 105 | TNFa_cDCs_pMAPKAPK2 | 9 |
| 106 | TNFa_cDCs_pP38 | 9 |
| 107 | TNFa_cDCs_pTBK1 | 9 |
| 108 | TNFa_NK.Cells_pCREB | 9 |
| 109 | TNFa_NK.Cells_pP38 | 9 |
| 110 | IFNa2_CD4.T.cells_pSTAT4 | 10 |
| 111 | IFNa2_CD4.T.cells_pSTAT5 | 10 |
| 112 | IFNa2_CD4.T.cells_pSTAT6 | 10 |
| 113 | IFNa2_CD8.T.cells_pSTAT4 | 10 |
| 114 | IFNa2_CD8.T.cells_pSTAT5 | 10 |

| | | |
|---|---|---|
| 115 | IFNa2_cDCs_pSTAT5 | 10 |
| 116 | IFNa2_cDCs_pSTAT6 | 10 |
| 117 | IFNa2_NK.Cells_pSTAT4 | 10 |
| 118 | IFNb_basophils_pSTAT6 | 10 |
| 119 | IFNb_CD4.T.cells_pSTAT4 | 10 |
| 120 | IFNb_CD4.T.cells_pSTAT5 | 10 |
| 121 | IFNb_CD4.T.cells_pSTAT6 | 10 |
| 122 | IFNb_CD8.T.cells_pSTAT4 | 10 |
| 123 | IFNb_CD8.T.cells_pSTAT5 | 10 |
| 124 | IFNb_CD8.T.cells_pSTAT6 | 10 |
| 125 | IFNb_NK.Cells_pSTAT4 | 10 |
| 126 | IFNb_NK.Cells_pSTAT5 | 10 |
| 127 | IFNb_NK.Cells_pSTAT6 | 10 |
| 128 | IL.4_B.Cells_pSTAT6 | 10 |
| 129 | IL.4_CD14..Monocytes_pSTAT6 | 10 |
| 130 | IL.4_CD16..Monocytes_pSTAT6 | 10 |
| 131 | IL.4_CD4.T.cells_pSTAT6 | 10 |
| 132 | IL.4_CD8.T.cells_pSTAT6 | 10 |
| 133 | IL.4_cDCs_pSTAT6 | 10 |
| 134 | IL.4_Neutrophils_pSTAT6 | 10 |
| 135 | IL.4_NK.Cells_pSTAT6 | 10 |
| 136 | IL.6_CD4.T.cells_pSTAT5 | 10 |
| 137 | CD40L_B.Cells_pCREB | 11 |
| 138 | CD40L_B.Cells_pErk1.2 | 11 |
| 139 | CD40L_B.Cells_pMAPKAPK2 | 11 |
| 140 | CD40L_B.Cells_pP38 | 11 |
| 141 | CD40L_B.Cells_pTBK1 | 11 |
| 142 | GM.CSF_CD14..Monocytes_pMAPKAPK2 | 11 |
| 143 | GM.CSF_CD16..Monocytes_pMAPKAPK2 | 11 |
| 144 | GM.CSF_cDCs_pP38 | 11 |
| 145 | GM.CSF_Neutrophils_pMAPKAPK2 | 11 |
| 146 | IFNg_cDCs_pSTAT5 | 11 |
| 147 | IL.4_B.Cells_pSTAT5 | 11 |
| 148 | IL.6_CD14..Monocytes_pSTAT3 | 11 |
| 149 | LPS_CD14..Monocytes_pMAPKAPK2 | 11 |
| 150 | LPS_CD14..Monocytes_pTBK1 | 11 |
| 151 | LPS_CD16..Monocytes_pErk1.2 | 11 |
| 152 | LPS_CD16..Monocytes_pMAPKAPK2 | 11 |
| 153 | LPS_CD16..Monocytes_pTBK1 | 11 |

| | | |
|---|---|---|
| 154 | LPS_cDCs_pErk1.2 | 11 |
| 155 | LPS_cDCs_pMAPKAPK2 | 11 |
| 156 | LPS_cDCs_pP38 | 11 |
| 157 | LPS_cDCs_pTBK1 | 11 |
| 158 | LPS_Neutrophils_pMAPKAPK2 | 11 |
| 159 | LPS_Neutrophils_pP38 | 11 |
| 160 | PMAIono_CD16..Monocytes_pErk1.2 | 11 |
| 161 | PMAIono_cDCs_pCREB | 11 |
| 162 | PMAIono_cDCs_pErk1.2 | 11 |
| 163 | R848_CD14..Monocytes_pMAPKAPK2 | 11 |
| 164 | R848_CD14..Monocytes_pTBK1 | 11 |
| 165 | R848_CD16..Monocytes_pErk1.2 | 11 |
| 166 | R848_CD16..Monocytes_pMAPKAPK2 | 11 |
| 167 | R848_CD16..Monocytes_pTBK1 | 11 |
| 168 | R848_cDCs_pErk1.2 | 11 |
| 169 | R848_cDCs_pMAPKAPK2 | 11 |
| 170 | R848_cDCs_pP38 | 11 |
| 171 | R848_cDCs_pTBK1 | 11 |
| 172 | R848_Neutrophils_pMAPKAPK2 | 11 |
| 173 | TNFa_Neutrophils_pMAPKAPK2 | 11 |
| 174 | Anthrax_CD14..Monocytes_IkBa | Unassigned |
| 175 | Anthrax_CD14..Monocytes_pTBK1 | Unassigned |
| 176 | Anthrax_CD16..Monocytes_pTBK1 | Unassigned |
| 177 | Anthrax_cDCs_IkBa | Unassigned |
| 178 | Anthrax_cDCs_pErk1.2 | Unassigned |
| 179 | GM.CSF_basophils_pMAPKAPK2 | Unassigned |
| 180 | GM.CSF_basophils_pSTAT5 | Unassigned |
| 181 | IFNa2_basophils_pSTAT5 | Unassigned |
| 182 | IFNb_basophils_pSTAT5 | Unassigned |
| 183 | IFNg_basophils_pSTAT5 | Unassigned |
| 184 | IL.12_NK.Cells_pSTAT4 | Unassigned |
| 185 | IL.2_NK.Cells_pErk1.2 | Unassigned |
| 186 | IL.2_NK.Cells_pSTAT6 | Unassigned |
| 187 | LPS_basophils_pP38 | Unassigned |
| 188 | LPS_CD14..Monocytes_p4E.BP1 | Unassigned |
| 189 | LPS_CD16..Monocytes_p4E.BP1 | Unassigned |
| 190 | LPS_Neutrophils_pTBK1 | Unassigned |
| 191 | PMAIono_basophils_pCREB | Unassigned |
| 192 | PMAIono_CD14..Monocytes_p4E.BP1 | Unassigned |

| | | |
|---|---|---|
| 193 | PMAIono_CD16..Monocytes_pCREB | Unassigned |
| 194 | PMAIono_cDCs_IkBa | Unassigned |
| 195 | PMAIono_cDCs_p4E.BP1 | Unassigned |
| 196 | PMAIono_cDCs_pSTAT1 | Unassigned |
| 197 | PMAIono_Neutrophils_pCREB | Unassigned |
| 198 | R848_Neutrophils_pTBK1 | Unassigned |
| 199 | TNFa_Neutrophils_pTBK1 | Unassigned |

Supplementary Table 6: Reduced set of immune features

| **Signaling proteins:** | **Conditions:** | **Proposed panel:** |
|---|---|---|
| p-STAT1 | Unstimulated control | CD45 |
| p-STAT5 | TNFa | CD3 |
| IkB | IFNa | CD4 |
| p-P38 | LPS | CD14 |
| | | p-STAT1 |
| **Cell types:** | | p-STAT5 |
| CD4+ T cells | | IkB |
| CD14+ Monocytes | | p-P38 |

Suggestion for an immune monitoring assay by traditional flow cytometry. The listed signaling proteins, cell types, and conditions provide surrogate markers for 10 of the 11 modules identified in our study and can be detected using an 8-parameter flow cytometry panel.

Supplementary Table 7: Selected features and coefficients for lasso model of sex

| Lasso model features | Coefficient |
|---|---|
| (Intercept) | 0.228368 |
| Anthrax_CD14..Monocytes_pTBK1 | 0.230217 |
| GM-CSF_CD16..Monocytes_pErk1.2 | -0.127964 |
| GM-CSF_Neutrophils_pErk1.2 | -0.109782 |
| IFNb_NK.Cells_pSTAT1 | 0.079948 |
| LPS_CD14..Monocytes_pTBK1 | 0.120808 |
| LPS_CD16..Monocytes_pP38 | -0.098301 |
| LPS_Neutrophils_pTBK1 | 0.138752 |
| PMAIono_Neutrophils_pCREB | -1.729958 |
| R848_Neutrophils_pP38 | -0.115437 |

Supplementary Table 8: Selected features and coefficients for sparse group lasso model of sex

| | Immune Feature | Module | SGL_coefficients |
|---|---|---|---|
| 1 | GM.CSF_basophils_pMAPKAPK2 | Unassigned | 1.702416025 |
| 2 | IFNa2_basophils_pSTAT5 | Unassigned | -1.034555697 |
| 3 | IFNb_basophils_pSTAT5 | Unassigned | -1.398076346 |
| 4 | IL.2_NK.Cells_pErk1.2 | Unassigned | 1.10622563 |
| 5 | IL.2_NK.Cells_pSTAT6 | Unassigned | 0.877075155 |
| 6 | LPS_CD14..Monocytes_p4E.BP1 | Unassigned | -0.560422876 |
| 7 | PMAIono_Neutrophils_pCREB | Unassigned | -5.924249754 |
| 8 | IFNb_CD8.T.cells_pSTAT1 | 1 | 1.378755128 |
| 9 | IFNb_NK.Cells_pSTAT1 | 1 | 0.680746216 |
| 10 | PMAIono_B.Cells_pMAPKAPK2 | 3 | 2.497055833 |
| 11 | PMAIono_CD8.T.cells_pCREB | 3 | 1.321243427 |
| 12 | CD40L_B.Cells_IkBa | 4 | -0.554788841 |
| 13 | Anthrax_basophils_pP38 | 5 | 4.066747714 |
| 14 | Anthrax_CD14..Monocytes_pP38 | 5 | 0.441602087 |
| 15 | GM.CSF_basophils_pCREB | 7 | -1.571687267 |
| 16 | GM.CSF_basophils_pP38 | 7 | -0.278224162 |
| 17 | GM.CSF_CD16..Monocytes_pErk1.2 | 7 | -0.58093994 |
| 18 | IL.6_CD14..Monocytes_pErk1.2 | 7 | -0.891622738 |
| 19 | LPS_CD16..Monocytes_pP38 | 7 | -3.033962639 |
| 20 | R848_Neutrophils_pP38 | 7 | -0.655153628 |
| 21 | TNFa_Neutrophils_pP38 | 7 | -3.878806731 |
| 22 | GM.CSF_Neutrophils_pSTAT5 | 8 | -1.484664026 |
| 23 | IFNb_Neutrophils_pSTAT5 | 8 | -0.348565349 |
| 24 | IL.2_CD4.T.cells_pSTAT5 | 8 | 6.948497596 |
| 25 | IFNa2_CD4.T.cells_pSTAT6 | 10 | 0.752503244 |
| 26 | IFNa2_cDCs_pSTAT5 | 10 | 0.598627434 |
| 27 | IL.4_B.Cells_pSTAT6 | 10 | 0.0091394 |
| 28 | IL.4_CD8.T.cells_pSTAT6 | 10 | 0.826159305 |
| 29 | IL.4_NK.Cells_pSTAT6 | 10 | 0.240081579 |
| 30 | CD40L_B.Cells_pErk1.2 | 11 | -0.007173256 |
| 31 | GM.CSF_CD14..Monocytes_pMAPKAPK2 | 11 | 1.41269069 |
| 32 | LPS_CD14..Monocytes_pTBK1 | 11 | 3.070859589 |
| 33 | PMAIono_CD16..Monocytes_pErk1.2 | 11 | -4.043346664 |
| 34 | R848_CD14..Monocytes_pMAPKAPK2 | 11 | 0.016602686 |
| 35 | R848_CD14..Monocytes_pTBK1 | 11 | 0.01868397 |

Supplementary Table 9: Selected features and coefficients for lasso model on module scores to predict sex

| Selected features | Coefficient |
|---|---|
| (Intercept) | 0.608973 |
| Module 1 score | 1.091937 |
| Module 5 score | 0.237815 |
| Module 7 score | -2.907148 |

Supplementary Table 10: Selected features and coefficients for lasso model of age

| Immune Feature | Model coefficient |
|---|---|
| (Intercept) | 41.49003261 |
| IFNb_CD8 T cells pSTAT1* | -20.84989452 |
| IL-2 NK Cells pErk1/2* | -19.13239344 |
| TNFa Neutrophils pMAPKAPK2* | 17.90804143 |
| IFNb CD4 T cells pSTAT4* | 15.31219672 |
| IFNb NK Cells pSTAT5* | -4.263942858 |
| R848 CD16 Monocytes pP38* | -3.096462489 |
| IFNa2 cDCs pSTAT1* | 2.932547271 |
| CD40L B Cells pP38* | -2.449038882 |
| IL-2 NK Cells pSTAT6* | -1.343930478 |
| Anthrax CD14 Monocytes pP38* | -0.763257612 |
| R848 cDCs pErk1/2* | 0.326302732 |
| GM-CSF cDCs pCREB | 0.08694423 |

Features selected and their coefficients from the lasso model regressing on age. Features marked with an asterisk were also selected in the sparse group lasso model.

Supplementary Figure 1:

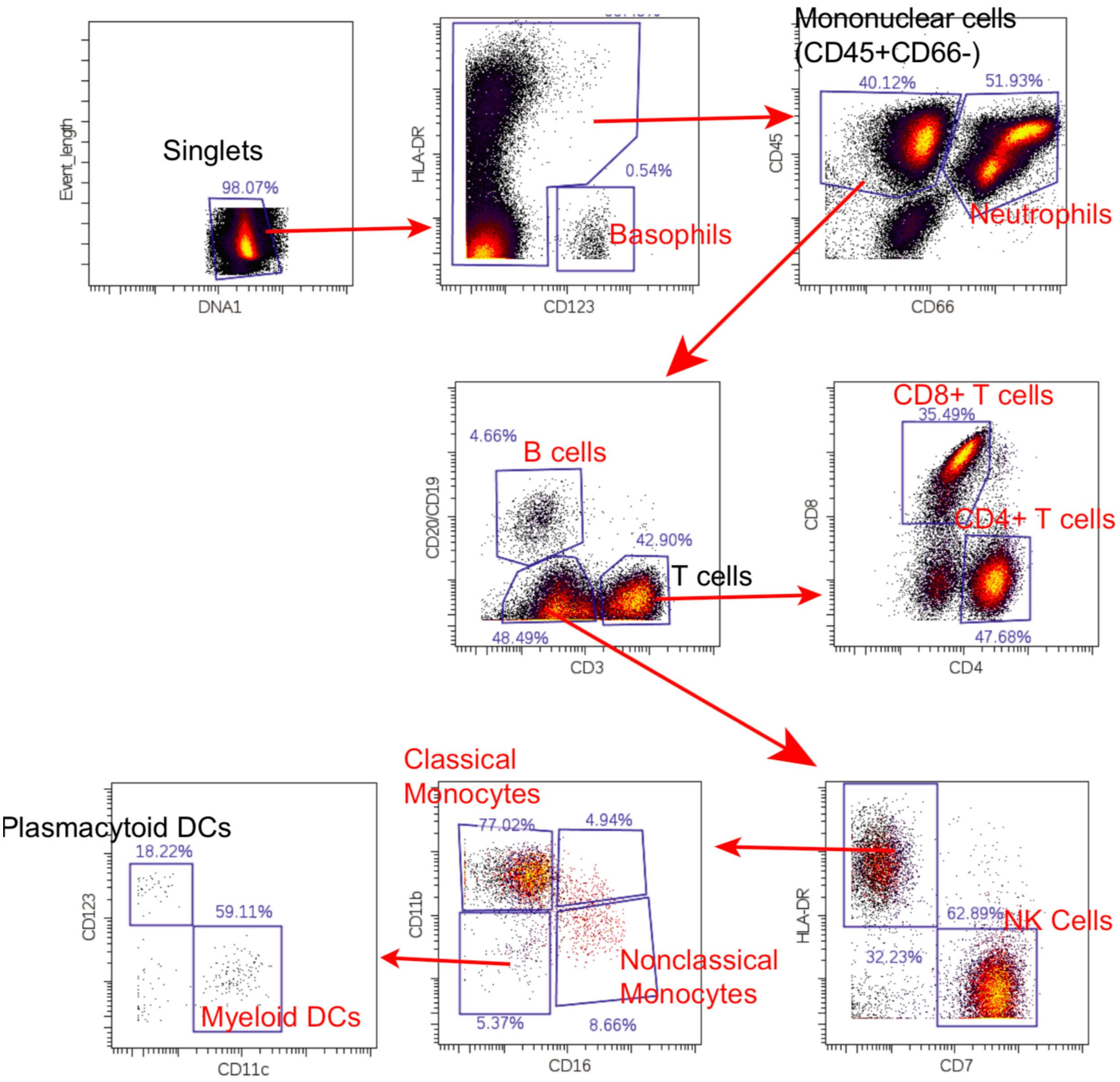


Manual gating strategy for all data (one representative donor shown). Populations used for analysis shown in red.

Supplementary Figure 2: Surrogate marker performance

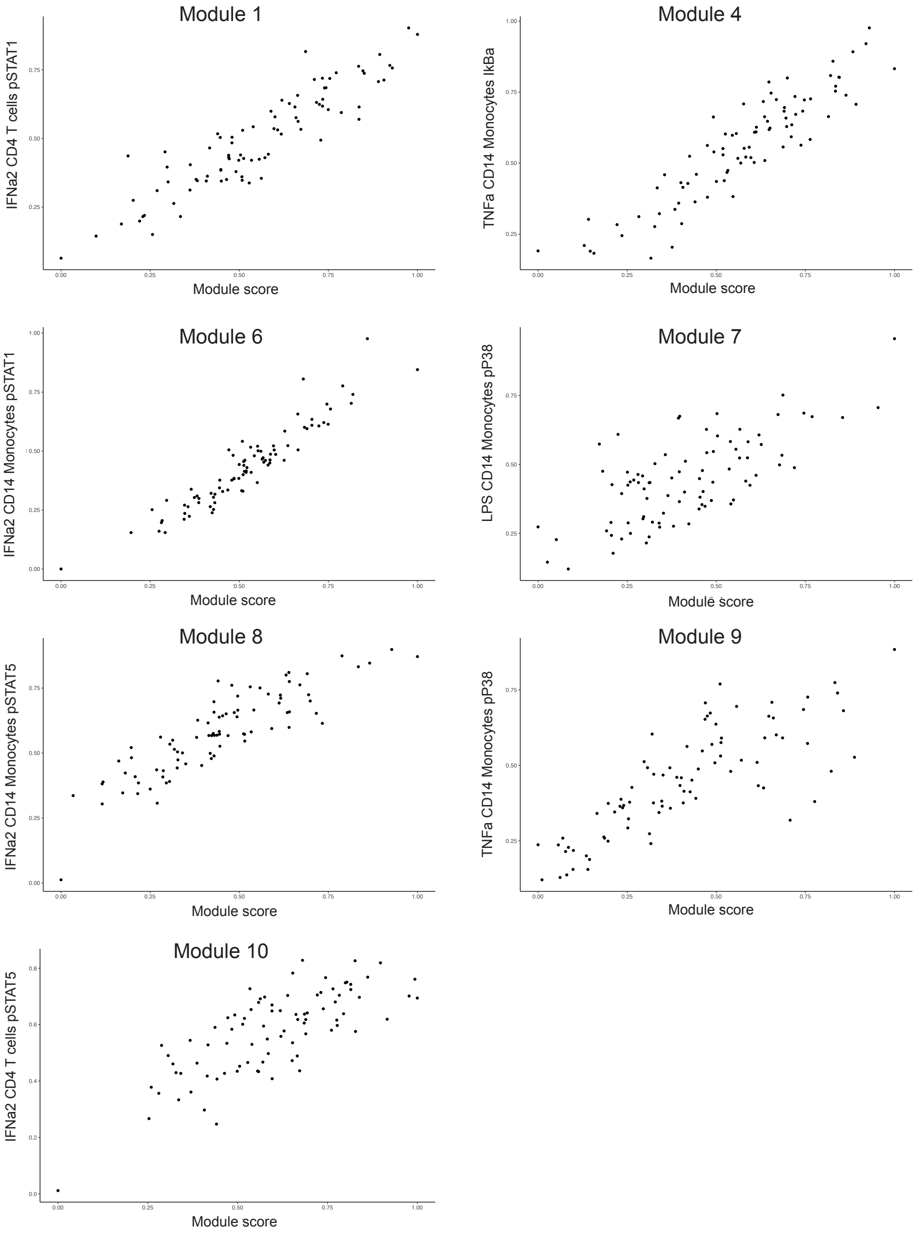


Comparison of individual's surrogate markers from a reduced feature set relative to their module scores using full data set. Value on y-axis is value of immune feature normalized to the range of that feature. Spearman rank correlation are as follows for modules 1, 4, 6, 7, 8, 9, 10: 0.90, 0.91, 0.94, 0.63, 0.88, 0.83, 0.74, respectively.

Supplementary Figure 3: Adjacency networks by age

Adjacency matrix in younger individuals:

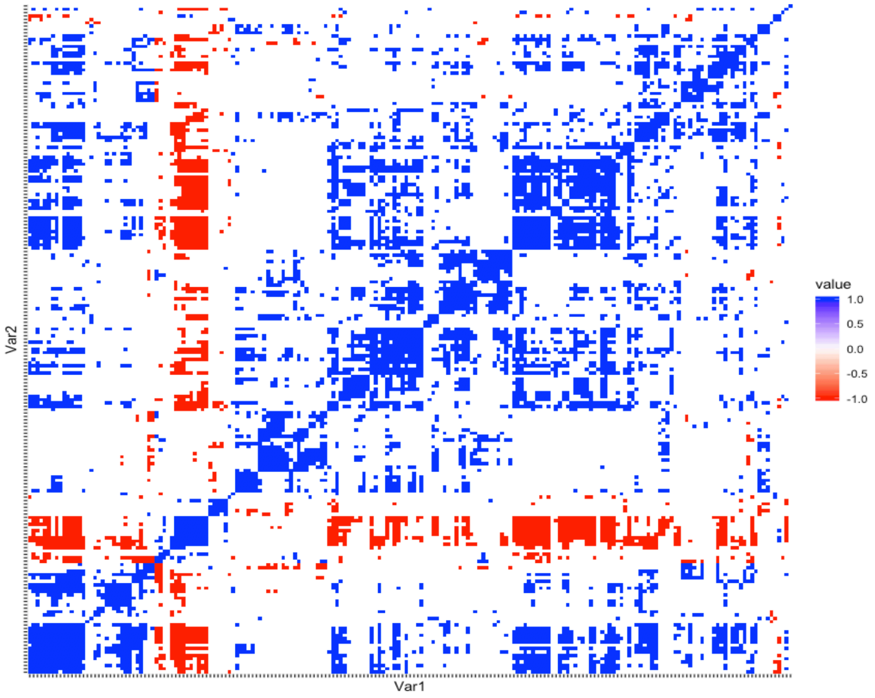


Adjacency matrix in older individuals:

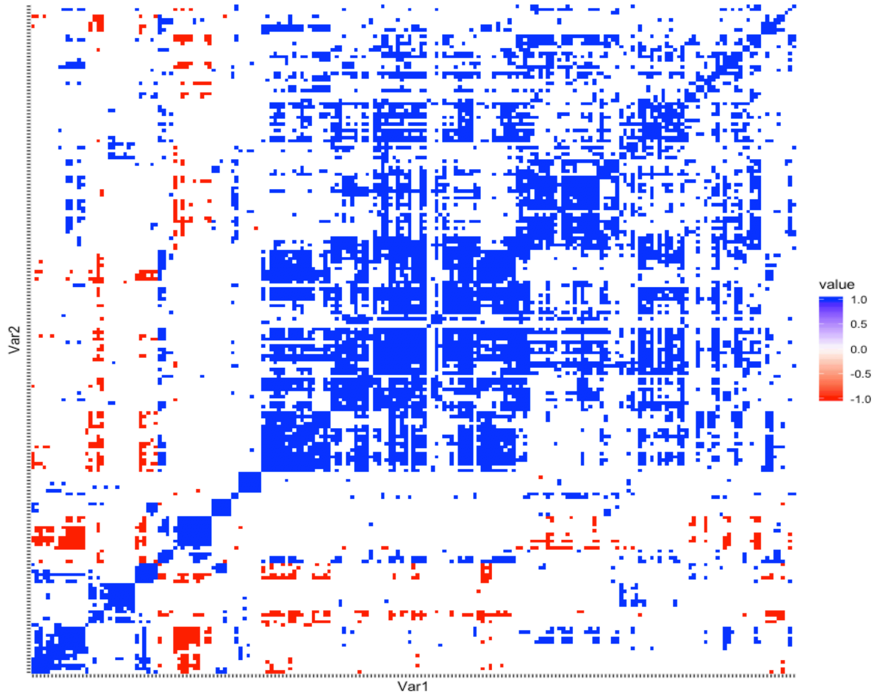


For gender balance, 10 males and 10 females were sampled for the younger individuals (≤ 30), 10 males and 10 females were sampled for the older individuals (≥ 45). Order was preserved from Figure 2.

Supplementary Figure 4: Correlation of module scores

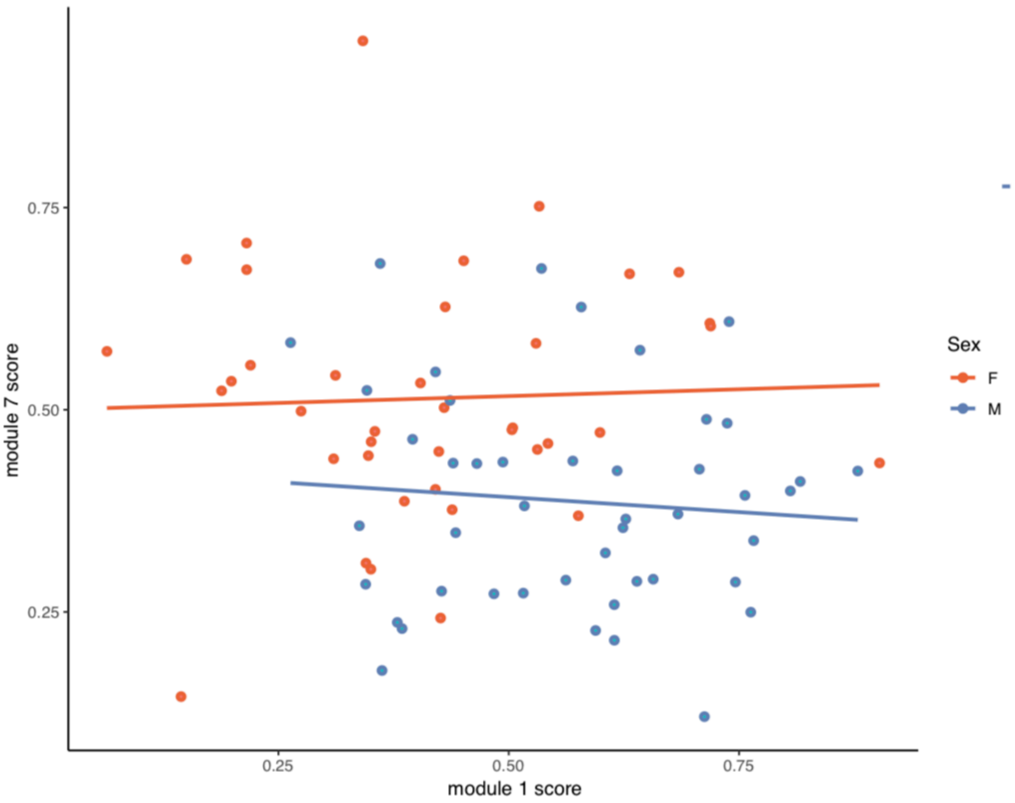


Module 1 scores do not correlate with module 7 scores within the sexes.

Supplementary Figure 5: Module score comparison between sexes

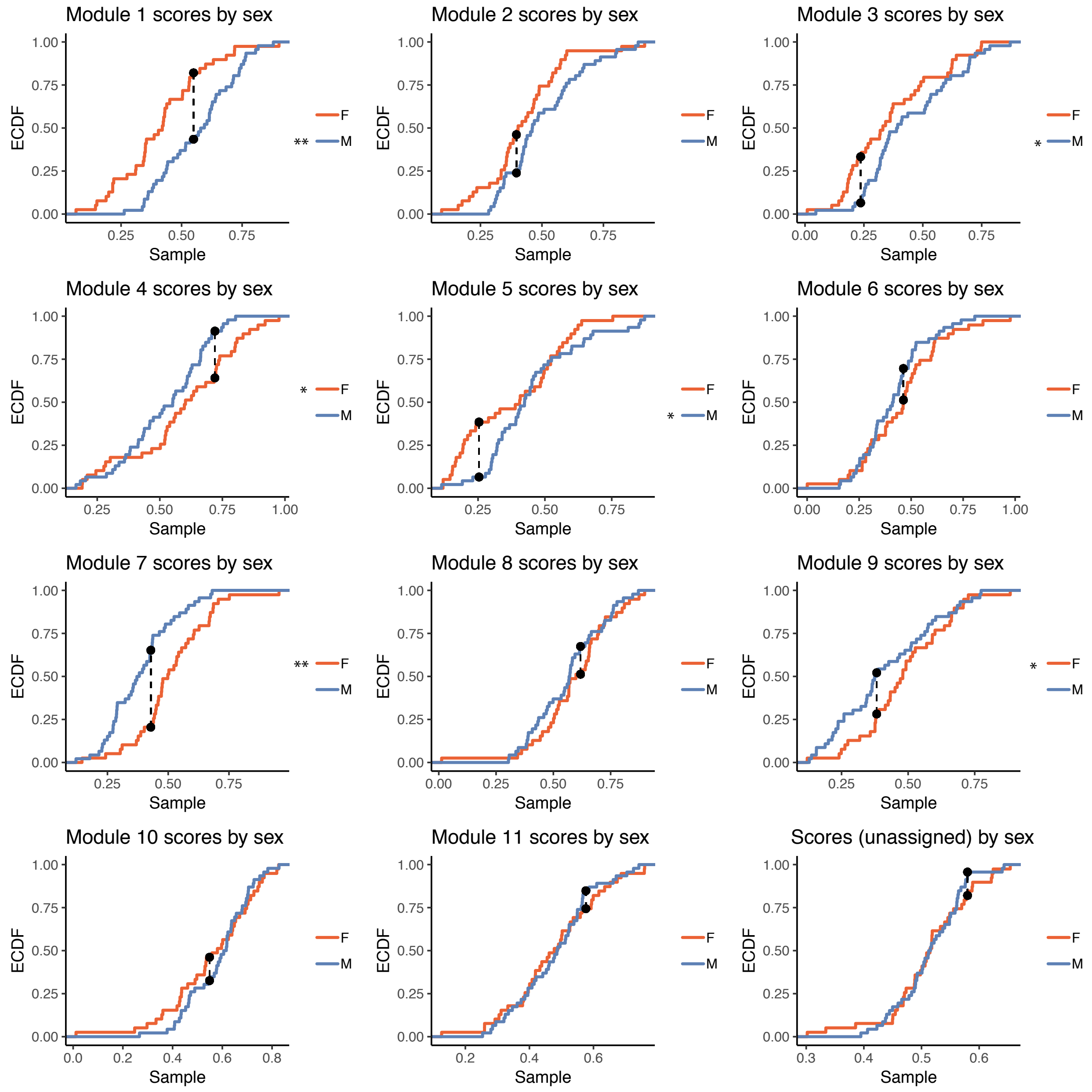


Comparison of module scores between sexes. Distributions marked with ** are significantly different when correcting for multiple hypothesis testing (Kolmogorov-Smirinov test, adjusted p-value < 0.05, h = 12). To show trends, all are modules are shown, those marked with * are significant without correcting for multiple hypothesis testing (Kolmogorov-Smirinov test, unadjusted p-value < 0.05).

Supplementary Figure 6: Sex differences in frequency data

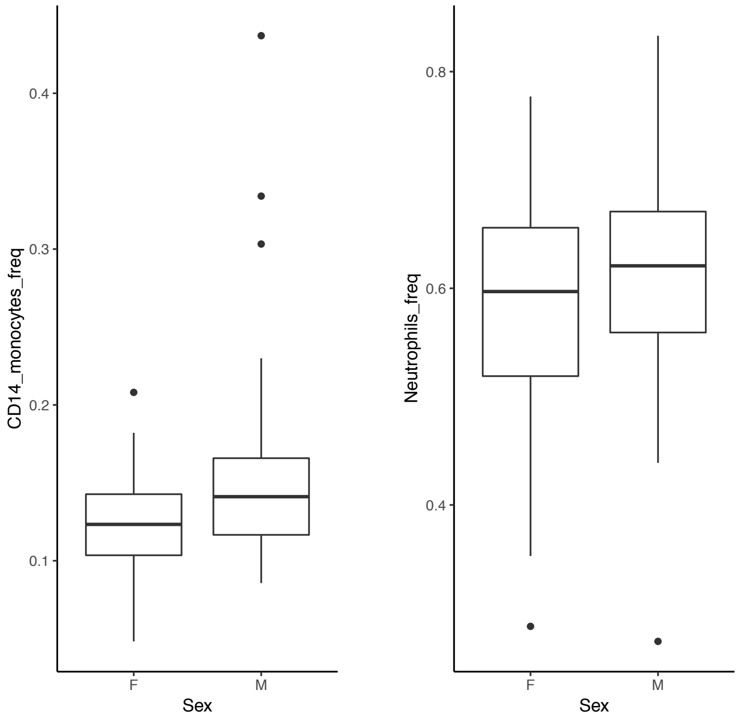


CD14+ monocyte frequencies (of mononuclear cells) and neutrophil frequencies (of singlets) in samples from male and female donors. Frequencies are higher in samples from male donors (Significance analysis of microarrays, FDR < .01).

Supplementary Figure 7: Tuning alpha parameter for elastic net

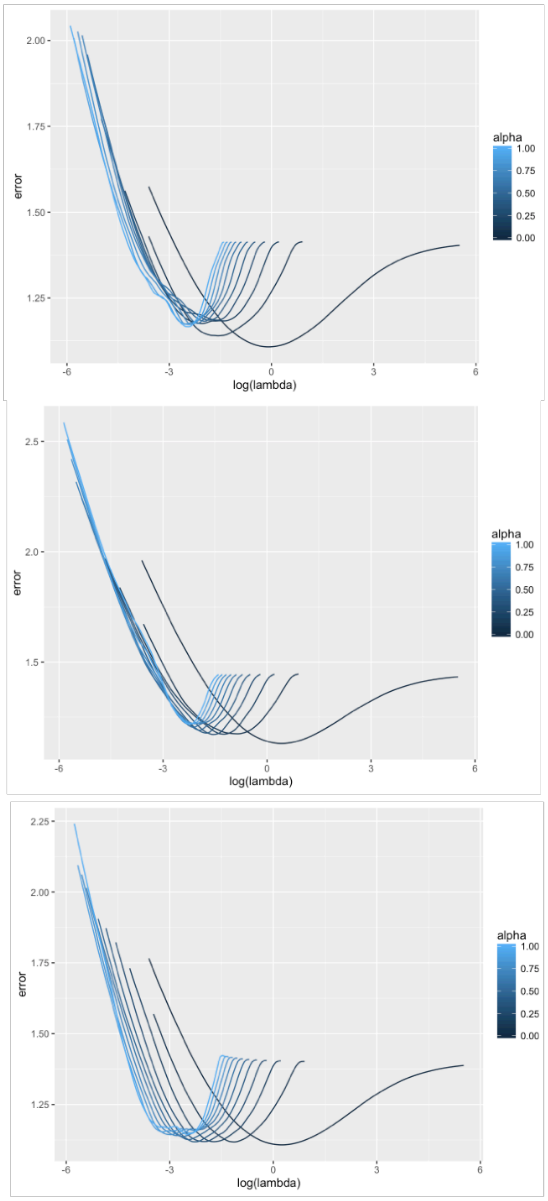


Selection of alpha for elastic net modeling of sex differences. Three iterations of assigning k-groups for cross-validation are shown. Error is lowest in all three cases using alpha = 0 (ridge model).
